# An allosteric switch regulates *Mycobacterium tuberculosis* ClpP1P2 protease function as established by cryo-EM and methyl-TROSY NMR

**DOI:** 10.1101/2019.12.11.873281

**Authors:** Siavash Vahidi, Zev A. Ripstein, Jordan B. Juravsky, Enrico Rennella, Alfred L. Goldberg, Anthony K. Mittermaier, John L. Rubinstein, Lewis E. Kay

## Abstract

The 300-kDa ClpP1P2 protease from *Mycobacterium tuberculosis* collaborates with the AAA+ (ATPases associated with a variety of cellular activities) unfoldases, ClpC1 and ClpX, to degrade substrate proteins. Unlike in other bacteria, all the components of the Clp system are essential for growth and virulence of mycobacteria, and their inhibitors show promise as novel antibiotics. MtClpP1P2 is unique in that it contains a pair of distinct ClpP1 and ClpP2 rings and also requires the presence of activator peptides, such as benzoyl-leucyl-leucine (Bz-LL), for function. Understanding the structural basis for this requirement has been elusive but is critical for the rational design and improvement of anti-TB therapeutics that target the Clp system. Here we present a combined biophysical and biochemical study to explore the structure-dynamics-function relationship in MtClpP1P2. Cryo-EM structures of apo and acyldepsipeptide-bound MtClpP1P2 explain their lack of activity by showing loss of a key β-sheet in a sequence known as the handle region that is critical for the proper formation of the catalytic triad. Methyl transverse relaxation-optimized spectroscopy (TROSY)-based NMR, cryo-EM, and biochemical assays show that upon binding Bz-LL or covalent inhibitors, MtClpP1P2 undergoes a conformational change from an inactive compact state to an active extended structure that can be explained by a modified Monod-Wyman-Changeux model. Our study establishes a critical role for the handle region as an on/off switch for function, and shows extensive allosteric interactions involving both intra- and inter-ring communication that regulate MtClpP1P2 activity and that can potentially be exploited by small molecules to target *M. tuberculosis*.

**Significance Statement:** The MtClpP1P2 protease is part of the essential protein degradation machinery that helps maintain protein homeostasis in *Mycobacterium tuberculosis*, the causative agent of TB. Antibiotics that selectively kill both dormant and growing drug-resistant populations of *M. tuberculosis* by disrupting MtClpP1P2 function have attracted recent attention. Here we characterize a switch that can control MtClpP1P2 activity through binding of small peptides, leading to a concerted conformational change that potentially can be exploited by drug molecules to interfere with MtClpP1P2 function. Overall, this work highlights the power of a combined NMR and cryo-EM approach to provide detailed insights into the structure-dynamics-function relationship of molecular machines critical to human health.

## Introduction

Tuberculosis (TB) is an infectious disease caused by *Mycobacterium tuberculosis* that typically affects the lungs of patients (1). Without proper treatment the mortality rate from TB is high, with approximately 1.5 million deaths per 10 million new cases each year (2). TB is thus the leading cause of mortality from a single infectious agent worldwide (2). Patients with compromised immune systems, such as patients treated with immunosuppressive drugs (3) or infected by HIV (4) are particularly vulnerable to infection. Drug-resistant strains of *M. tuberculosis* are increasingly common (2, 5) with one-half million cases of TB in 2017 resistant to rifampicin, the standard first-line drug therapeutic (2). Treatments for drug-resistant strains are complex, slow, expensive, and have severe side-effects (6–8). Notably, it is estimated that one quarter of the world’s population is infected with latent *M. tuberculosis* (9, 10). These alarming facts emphasize the urgent need to develop novel therapeutic strategies that are effective against drug-resistant strains of *M. tuberculosis*, which in turn requires an atomic level understanding of the key molecular components that are responsible for maintaining the viability of this disease-causing pathogen.

The ClpP protease is a major component of one of the main protein degradation systems in many bacteria (11–14). ClpP is essential for homeostasis, pathogenesis and virulence in several bacteria (12, 15, 16) and is considered a novel drug target for antibiotics that disrupt enzyme function as their mechanism of action (17–21). For example, cyclic acyldepsipeptides (ADEPs) cause unregulated activity in most bacterial ClpPs leading to the proteolysis of proteins necessary for survival, and have proven to be an effective bactericide against antibiotic-resistant and persistent subpopulations of *Staphylococcus aureus* (22–25). In *M. tuberculosis*, ClpP1P2 (hereafter referred to as MtClpP1P2) is essential for growth and virulence (26, 27), and requires the binding of essential AAA+ unfoldases, ClpX or ClpC1, that use the energy of ATP to unfold and translocate substrates into its catalytic chamber for degradation (28). Lassomycin (29), ecumicin (30), and rufomycin (31) are promising antibiotics that selectively kill both dormant and growing drug-resistant populations of *M. tuberculosis* by binding to ClpC1 and decoupling ATP-dependant protein unfolding from proteolysis. Similarly, ADEPs also kill *M. tuberculosis* by preventing the binding of AAA+ regulatory unfoldases to MtClpP1P2 (32). MtClpP1P2 reacts with standard inhibitors of serine proteases, such as chloromethyl ketones, which modify the serine and histidine residues at enzyme active sites (33). Peptide boronates have been shown to also directly engage MtClpP1P2 active sites, causing inhibition at low micromolar concentrations and preventing growth of *M. tuberculosis* (34–36). More recently Cediranib, an anti-cancer drug, was proposed as a novel non-covalent inhibitor of MtClpP1P2 (37).

Most bacteria possess a single *clpP* gene, giving rise to a structure comprising a pair of heptameric rings that are arranged coaxially to form a homotetradecameric barrel-like protein complex enclosing fourteen Asp-His-Ser catalytic triads (11, 12, 38). Each of the identical protomers consists of an N-terminal domain that forms gated narrow pores on the apical surface of the barrel, a head domain that generates the main body of the ClpP barrel, and a handle region comprising a helix and a β-sheet that mediate ClpP ring-ring interactions (11, 38–40). In addition, the handle provides crucial contacts that align the catalytic triad and generates a binding grove for substrate polypeptides (41). Opening of the pores that allow substrate translocation is tightly regulated by AAA+ regulators (42, 43).

Actinobacteria, the phylum to which mycobacteria such as *M. tuberculosis* belong, are unique in that they contain two *clpP* genes, *clpP1* and *clpP2*, that encode for MtClpP1 and MtClpP2, respectively (44, 45). Initial structure-function studies concluded that MtClpP1 and MtClpP2 were separate enzymes that in isolation formed mixtures of homo-heptameric and homo-tetradecameric complexes that lacked activity (46, 47). Later, the specific catalytically functional form of the enzyme was elucidated upon the addition of small-molecule activators, such as benzoyl-leucyl-leucine (Bz-LL), that promote the formation of heterocomplexes comprising one copy of each of the MtClpP1 and MtClpP2 rings, converting the inactive enzyme to a catalytic form upon binding (33). Recent studies (48) showed that mixing of MtClpP1 and MtClpP2 result in MtClpP1P2 complexes, even in the absence of activators. Unlike for other ClpPs, the addition of ADEP alone does not activate MtClpP1P2 (32, 49). The requirement for activators appears to be a unique feature of actinobacteria, as other species that contain two ClpP proteoforms, such as *Listeria monocytogenes* (50, 51) and *Chlamydia trachomatis* (52), are functional even in their absence. X-ray structures of MtClpP1P2 crystalized in the presence of Bz-LL show that the overall architecture and fold of each ClpP protomer is indistinguishable from what is found in other bacterial ClpPs that retain activity in the apo form (53). Paradoxically, dipeptide activators occupy all fourteen MtClpP1P2 catalytic sites in these structures, with activity arising presumably when the activators are displaced by substrate. Another crystal structure of MtClpP1P2 that includes ADEP and an IL-dipeptide activator confirms the full occupancy of peptide activator in all catalytic sites (49). In this structure seven ADEP molecules are bound exclusively to MtClpP2 and interact with the hydrophobic pocket at the interface between two adjacent protomers. Currently there are no structures available for the inactive, apo- and ADEP-bound MtClpP1P2 complexes in the absence of activators and hence both the structural basis for the lack of activity and the mechanism of MtClpP1P2 activation by peptides remains unclear. The lack of activity is particularly intriguing given that other ClpPs are active in the apo conformation, and that their activity increases upon ADEP-binding.

Herein we present a combined biophysical and biochemical study, using methyl-TROSY based NMR spectroscopy as well as electron cryo-microscopy (cryo-EM), to explore the structure-dynamics-function relationship in MtClpP1P2. We show that MtClpP1 and MtClpP2 exist as heptamers in solution, and that upon mixing they readily form MtClpP1P2 tetradecamers, even in the absence of activators. NMR titration data quantify the interaction between MtClpP1 and MtClpP2, establishing that binding becomes stronger in the presence of the Bz-LL activating dipeptide. Structures of the inactive apo-MtClpP1P2 and ADEP-bound MtClpP1P2 complexes, determined by cryo-EM, explain their lack of activity by revealing the absence of a key β-sheet in the handle region that is present in all active ClpP conformations. Methyl-TROSY based spectra show that upon binding Bz-LL a significant conformational change occurs for MtClpP1P2 that corresponds to a conversion from an inactive state to an active structure. The NMR based Bz-LL titration data and extensive activity profiles as a function of both Bz-LL and substrate concentrations can be explained by a modified Monod-Wyman-Changeux (MWC) model that highlights the extensive allostery in this system. Our study establishes a critical role for the handle region, serving as an on/off switch for function, and a strong network of allosteric interactions involving both intra- and inter-ring communication that regulate activity of the MtClpP1P2 enzyme.

## Results

### Probing the oligomeric state of MtClpP1P2

MtClpP1 and MtClpP2, lacking their propeptide sequences as established from previous work (33, 48), were overexpressed in *E. coli* and purified as detailed in SI. Size exclusion chromatography (SEC) was used to probe the oligomeric forms of MtClpP1 and MtClpP2 in isolation and when mixed. MtClpP1 (residues 7-200) elutes at the same retention volume as a separately isolated R171A mutant of ClpP from *Staphylococcus aureus* (SaClpP) that is known to form a heptameric single ring (23) (0.5 mL injected at 175 µM monomer – Fig. 1A, green trace; black arrow denotes the elution position of R171A SaClpP). Increasing or decreasing the injected protein concentration three-fold, from 175 µM monomer to 525 or 58 µM, respectively, does not change the SEC peak elution volume and profile (Fig. S1A), suggesting that the single ring is stable. SEC of the mature form of MtClpP2 with native propeptide processing that removed the N-terminal 12 residues (residues 13-214) revealed a pair of peaks at retention volumes of 11 and 13 mL that correspond to assemblies which are either larger or approximately the same size as tetradecameric WT SaClpP, respectively (0.5 mL injected at 100 µM monomer - Fig. S1B - blue trace). Unfortunately, this construct is aggregation-prone and polydisperse. Inspired by previous work (48), a MtClpP2 construct with three additional residues removed from the N terminus, Δ15 MtClpP2 (16-214), was used in the present study (Fig. S1C). Δ15 MtClpP2 exhibits indistinguishable peptidase activity in MtClpP1P2 complexes relative to the original MtClpP2 protein (Fig. S1D), and millimolar concentration solutions are stable for at least several weeks at 4 °C. SEC analysis of Δ15 MtClpP2 (hereafter referred to as MtClpP2) showed a purely heptameric form that elutes at the same volume as MtClpP1 (0.5 mL injected at 175 µM – Fig. 1A, blue trace) and its oligomeric distribution is similarly insensitive to protein concentration (Fig. S1C). Mixing of MtClpP1 and MtClpP2 yields a larger oligomeric species, corresponding to a MtClpP1P2 complex (0.5 mL injected at 175 µM of each of MtClpP1 and MtClpP2 - Fig. 1A, purple trace) that elutes at the same volume as the well-characterized tetradecameric SaClpP (Fig. 1A, black trace). Notably, the original (13-214) MtClpP2 construct was also able to bind MtClpP1 (Fig. S1B-purple trace).

**Figure 1.**
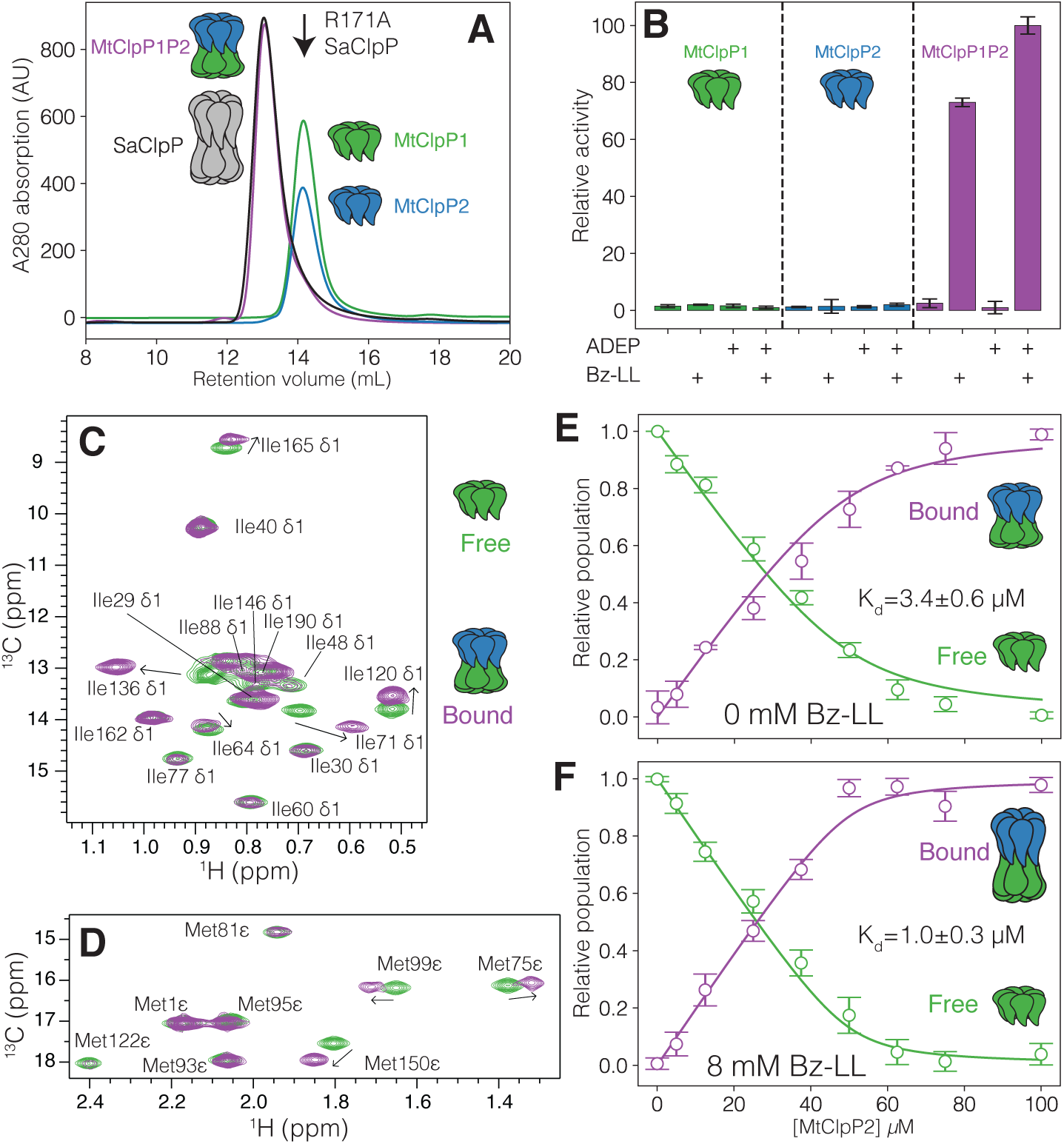
Characterization of the interaction between MtClpP1 and MtClpP2. (A) SEC profiles of isolated MtClpP1 (residues 7-200; green trace) and MtClpP2 (residues 16-214; blue trace) and a mixture containing equimolar amounts of the two (purple trace). The SEC elution profile of WT, tetradecameric SaClpP (black trace) and the elution volume of a R171A, heptameric mutant of SaClpP (black arrow) are included. In all cases 0.5 mL of protein at a concentration of 175 µM (monomer) was injected; (B) Peptidase activity assays, 40 °C, on isolated MtClpP1 (green bars) and MtClpP2 (blue bars) and mixtures containing equimolar amounts of the two (purple bars), performed in the presence or absence of Bz-LL (4 mM) and/or ADEP (25 µM) activators with 250 µM PKM-AMC used as substrate. In all cases the concentration of each of MtClpP1 and MtClpP2 was 1 µM (monomer concentration). Error bars correspond to one standard deviation based on three measurements. (C-D) Overlay of the (C) Ile and (D) Met regions of ^1^H-^13^C HMQC correlation maps of ILVM-labeled MtClpP1 (green contour) and a mixture containing 50 µM ILVM-labeled MtClpP1 and 100 µM [U-^2^H] MtClpP2 (purple contours), recorded at 40 °C, 18.8 T, with assignments. (E-F) Changes in the intensity of the apo (green circles) and bound peaks (purple circles) in a titration of [U-^2^H] MtClpP2 into 50 µM ILVM-labeled MtClpP1 in the (E) absence or (F) presence of 8 mM Bz-LL. Error bars correspond to one standard deviation measured across all peaks averaged and the solid lines are fits to a simple binding model (see SI). The measured dissociation constant is indicated on each plot.

### Characterization of the interaction between MtClpP1 and MtClpP2

Peptidase assays were performed to functionally characterize MtClpP1P2 by monitoring the fluorescence change resulting from cleavage of the fluorogenic substrate peptide N-Acetyl-Pro-Lys-Met-7-amino-4-methylcoumarin (PKM-AMC) (36). Notably, neither MtClpP1 or MtClpP2 in isolation was active (Fig. 1B – green and blue), with both MtClpP1, MtClpP2, and Bz-LL all required for activity (Fig. 1B – 2nd purple bar). The addition of ADEP to this mixture results in further activation (Fig. 1B – 4th purple bar). Unlike ClpPs from other organisms, MtClpP1P2 is not activated by ADEP alone (Fig. 1B – 3rd purple bar). These assays establish that the MtClpP1 and MtClpP2 constructs used in this work have similar functional properties as those found in previous reports (33, 48, 49, 53).

In order to characterize the binding between MtClpP1 and MtClpP2 more quantitatively we used methyl-TROSY NMR spectroscopy that exploits dipolar cross-correlated spin-relaxation to significantly improve spectral sensitivity and resolution in high molecular weight protein complexes (54–56). This approach is optimal in concert with selective isotopic labeling of methyl groups (^13^CH_3_), such as those of Ile, Leu, Val, and Met, within an otherwise uniformly deuterated ([U-^2^H]) protein, where the prochiral methyls of Leu/Val are ^13^CH_3_/^12^CD_3_ (referred to hereafter as ILVM-labeling) (57). Here ^1^H-^13^C heteronuclear multiple-quantum coherence (HMQC) correlation maps are recorded that benefit from this methyl-TROSY effect (54). High-quality spectra were obtained for samples of isolated ILVM-labeled MtClpP1 rings, and when perdeuterated, and hence NMR-invisible MtClpP2, is added to form a MtClpP1P2 complex (Fig. 1C). Notably, spectral quality for the MtClpP1 heptamer is superior to that for the corresponding MtClpP2 ring and for this reason we focused on molecules where MtClpP1 was labeled. Resonance assignments of MtClpP1 ILVM-methyl probes were obtained through a combined approach involving single-point mutagenesis and analysis of nuclear Overhauser effect (NOE) datasets, aided by the availability of high-resolution X-ray crystal structures of MtClpP1 (46, 53). The Ile (Fig. 1C – green contours) and Met (Fig. 1D – green contours) regions of a 2D ^1^H-^13^C HMQC spectrum of ILVM-labeled MtClpP1 recorded at 40 °C and 18.8 T have been completely assigned (17 Ile and 8 Met residues). Corresponding spectral regions and stereospecific assignments for all 7 Val, and 14 out of 18 Leu methyl residues are shown in the Supporting Information (Figure S2 – green contours). The addition of [U-^2^H] MtClpP2 leads to the emergence of separate peaks for residues proximal to the ring-ring interface and the handle region (e.g., Ile71, Ile120, Ile165, Val145, Met75, Met99, and Met150) (Fig. 1C&D and Figure S2 – purple contours), corresponding to the formation of MtClpP1P2, with exchange between apo- (MtClpP1) and bound- (MtClpP1P2) states slow on the chemical shift timescale. An NMR titration was performed to obtain binding constants for the MtClpP1 and MtClpP2 interaction (Fig. 1E). For residues with well-resolved peak displacements upon binding of MtClpP2, intensities of apo (Fig. 1E – green) and bound peaks (Fig. 1E – purple) were averaged and fitted with a simple binding model (see SI). This analysis yielded a dissociation constant of 3.4 ± 0.6 µM in the absence of Bz-LL activator. The interaction between MtClpP1 and MtClpP2 is moderately strengthened in the presence of 8 mM Bz-LL, with a dissociation constant of 1.0 ± 0.3 µM.

### Cryo-EM structure of apo MtClpP1P2 explains its lack of activity

Several crystal structures of MtClpP1P2 bound to dipeptide activators have been reported showing that the complex adopts the active extended conformation (53), similar to the structures of other ClpPs that are functional without the addition of activating molecules (38, 39, 58, 59). The absence of a high-resolution structure for apo-MtClpP1P2 has precluded an understanding of the underlying reason for the lack of activity for *M. tuberculosis* ClpP. To address this question MtClpP1 and MtClpP2 heptamers were mixed and the resulting apo MtClpP1P2 complex was isolated using SEC and subjected to cryo-EM analysis. Following 2D classification (Fig. S3 A&B), 373064 particle images were used for a final refinement with C7 symmetry, achieving a resolution of 3.1 Å (Fig. S4 A&B). Further classification without enforcing symmetry did not yield additional conformations. The refined map shows a tetradecameric arrangement, comprising a pair of heptameric MtClpP1 and MtClpP2 rings (Fig. 2A), akin to the Bz-LL bound MtClpP1P2 structure that was solved to 3.0 Å by X-ray crystallography (53) (Fig. 2B – PDBID: 5DZK) and subsequently confirmed in the present study. Whereas the Bz-LL bound structure is in the active extended conformation, the apo form appears to adopt the “compact” state, characterized by a shorter barrel height (83 Å vs 87 Å) (Fig. 2A&B). Compression of the barrel is accompanied by disruption of the handle β-sheet (Fig. 2C&D), which in the active conformation connects pairs of protomers from opposite rings. In the apo structure the two strands move apart and become more flexible, as suggested by lower resolution for these strands relative to the rest of the protein complex (Fig. 2E, Fig. S5A, Fig. S6). This disorder/flexibility is particularly notable for MtClpP2. The handle region is critically important for function as it positions the catalytic triad for optimal activity and provides a binding groove for ClpP substrates (Fig. 2C&D). In both MtClpP1 and MtClpP2 rings, the catalytic His residue (His123 in MtClpP1) is located three amino-acids N-terminal to the start of the handle β-sheet. The disordered conformation of the handle region displaces His123 from its position in the active conformation, thereby preventing the proper orientation of the catalytic triad residues (Fig. 2F&G). A well-known requirement of ClpP activity is engagement of oligomeric sensors comprising a pair of Asp-Arg salt bridges that link protomers in opposite rings and couple oligomerization with catalysis by stabilizing the active extended conformation (23, 50, 60). In contrast to the case in the apo-MtClpP1P2 structure where the oligomeric sensors are disengaged (Fig. 2H), in the Bz-LL bound structure all seven Asp170 - Arg171 salt bridges connecting opposite rings are observed (Fig. 2I).

**Figure 2.**
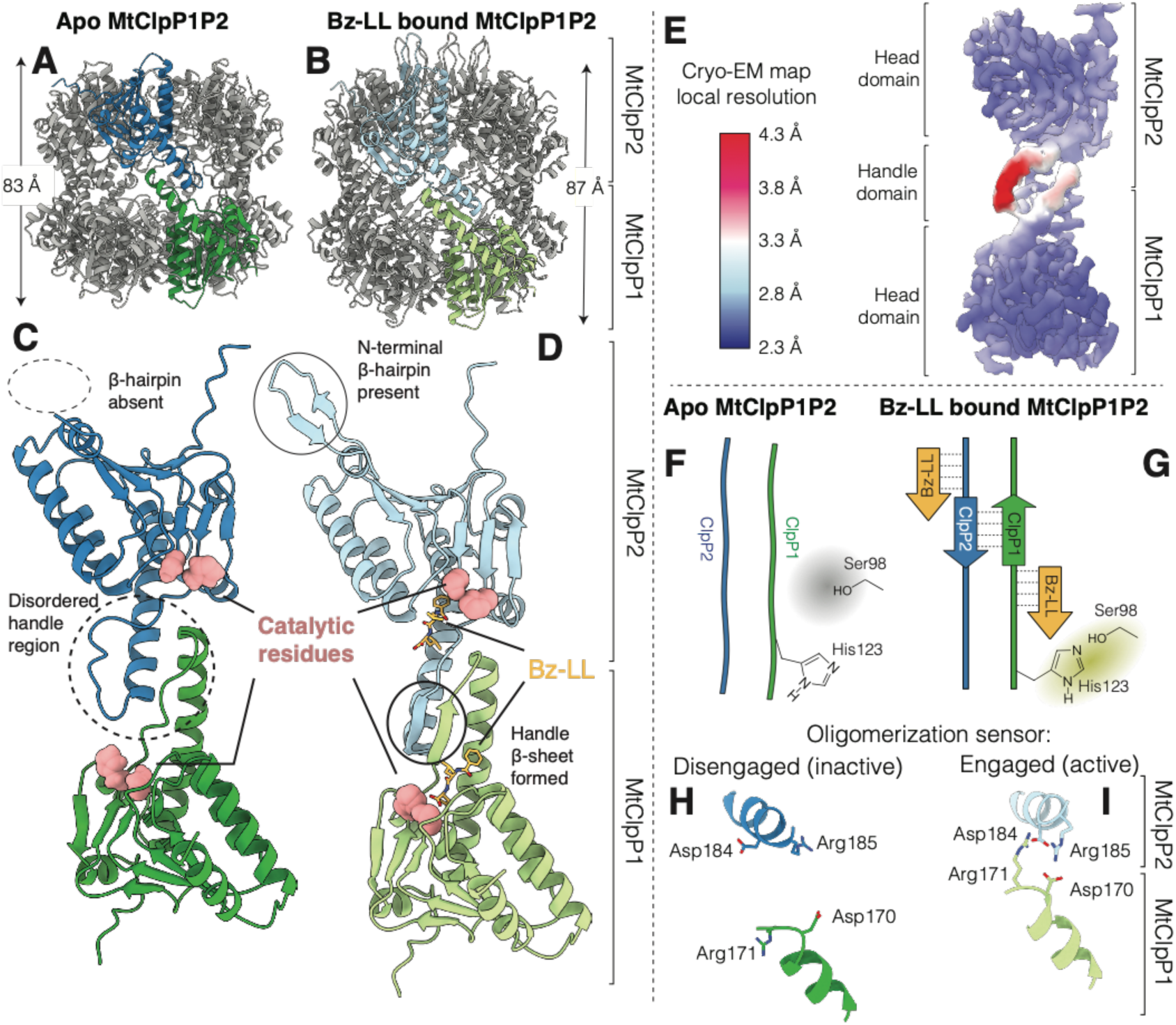
Structural basis for the lack of activity of apo MtClpP1P2. (A&C) Cryo-EM structure of apo MtClpP1P2 (this study), and (B&D) crystal structure of Bz-LL bound MtClpP1P2 obtained from ref. (53) (PDBID: 5DZK). In each case a pair of MtClpP1 (green) and MtClpP2 (blue) protomers are coloured to show the ring-ring interaction interface. The height of the ClpP barrel (excluding the N-terminal β- hairpins of MtClpP2) is denoted in each case. Major structural differences between apo and Bz-LL bound MtClpP1P2 are highlighted. (E) Cryo-EM density map of apo MtClpP1P2 at 3.1 Å resolution refined using C7 symmetry, with the distribution of local resolution colour-coded as indicated. (F&G) Schematic showing how the conformation of the handle β-sheets is affected by Bz-LL binding and its influence on positioning of the catalytic His. (H&I) Status of the oligomerization sensor in the apo and Bz-LL-bound forms of MtClpP1P2. Side chains of residues involved in salt-bridge formation are indicated.

### Structural basis for inactivity of ADEP-bound MtClpP12

The assays shown in Figure 1B establish that apo-ClpP1P2 is not catalytically active, with the structural basis for this lack of activity explained by the model of the apo-MtClpP1P2 complex showing a disordered handle region. Given that binding of a variety of different ADEPs fails to activate MtClpP1P2, including ADEP-7 assayed here, as well as six different analogues studied elsewhere (49), we postulated that this class of activator would not be able to reorganize the handle into a catalytically competent form. In order to test this hypothesis, we calculated a 3.1 Å resolution cryo-EM map of MtClpP1P2 bound to ADEP (Fig. S4 C&D). After 2D classification, 192430 particle images were used in a final refinement step with C7 symmetry enforced (Fig. S3 C&D). Further classification without enforcing symmetry did not reveal additional conformations. As in the apo structure, the height of the barrel is 83 Å (excluding N-terminal gates) (Fig. 3A). There is clear density for seven ADEP molecules at the interface of the MtClpP2 protomers (Fig. 3A - inset), with no ADEP bound to MtClpP1 (Fig. 3B&C), consistent with biochemical evidence (48) and published crystal structures of MtClpP1P2 bound to both ADEP and dipeptide activator (49). Compared to apo-MtClpP1P2, ADEP binding causes a striking rigidification of the MtClpP2 N-terminal β-hairpins (Fig. 3C). Importantly, the handle region remains disordered (Fig. 3C, Fig. S5B) and the oligomeric sensor is disengaged (Fig. 3D). Apart from the conformational changes that ADEP binding introduces to MtClpP2, the overall fold and conformation of MtClpP1 resembles that of the apo form, with a root-mean-square deviation (RMSD) between MtClpP1 rings of only 0.29 Å (Fig. 3B). Interestingly, activity response curves using PKM-AMC as substrate, for which only MtClpP1 has significant activity (Fig. S7), show a shift to lower Bz-LL values upon the addition of ADEP (Fig. 3E). This observation is consistent with an inter-ring allosteric communication originating from the ADEP binding site on the apical surface of MtClpP2 that propagates to the active sites of MtClpP1.

**Figure 3.**
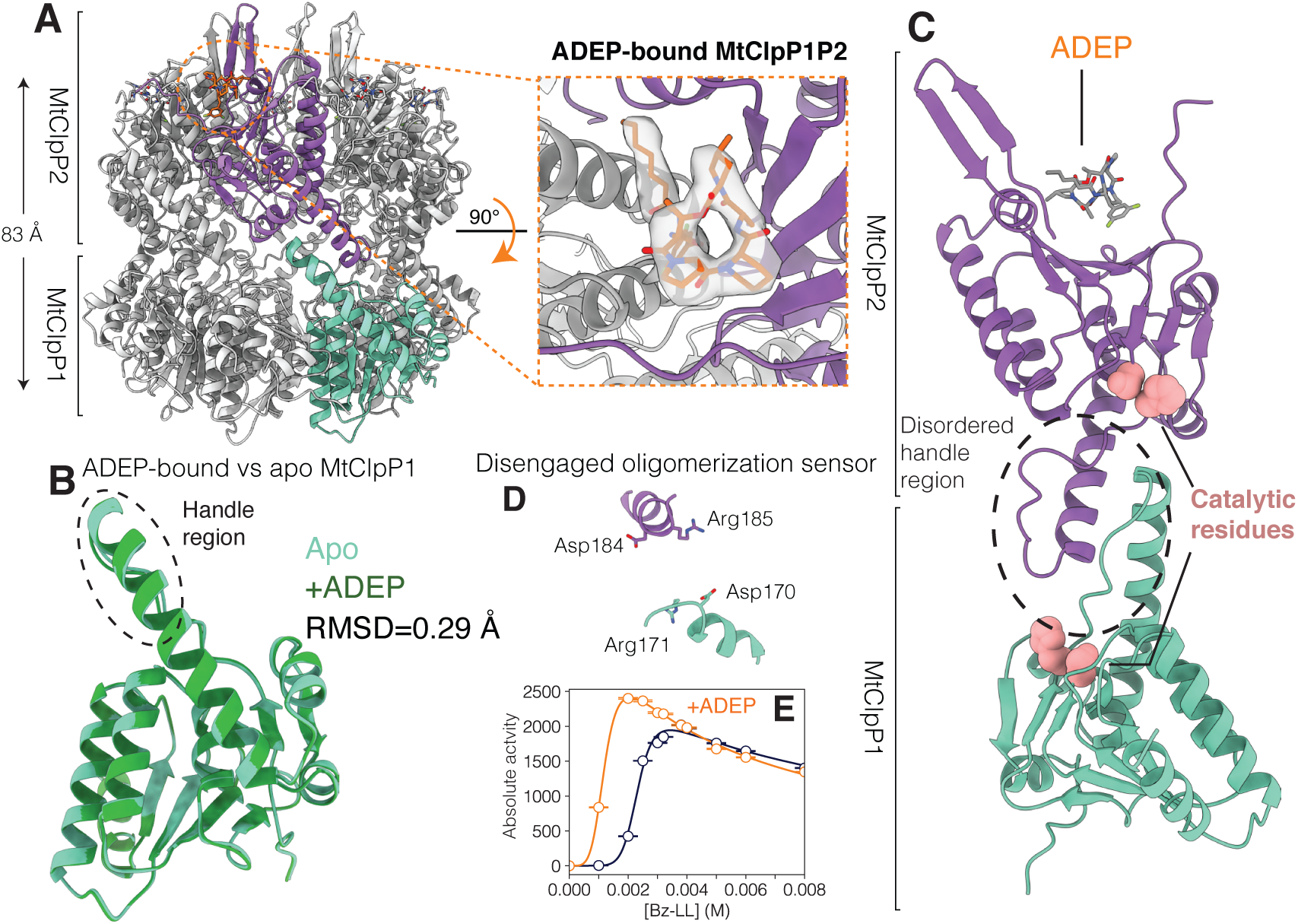
Structural basis for the lack of activity of ADEP-bound MtClpP1P2. (A) Side view of a cryo-EM structure of MtClpP1P2 bound to ADEP only. The height of the MtClpP1P2 barrel (excluding the N-terminal β-hairpins of MtClpP2) is denoted. Inset shows top view of the MtClpP1P2 complex looking down into the ADEP binding pocket at the interface of a pair of adjacent MtClpP2 protomers. The cryo-EM density corresponding to bound ADEP is displayed. (B) RMSD and overlay of single MtClpP1 protomers from MtClpP1P2 structures in the apo (light green) and the ADEP-bound (dark green) forms. Notably, the handle region (circled) remains disordered in the ADEP-bound form as in the apo structure. (C) Magnified view of a pair of opposite MtClpP1 (cyan) and MtClpP2 (purple) protomers, shown to highlight the disordered handle region and its position relative to the catalytic residues and the bound ADEP molecule. (D) Status of the oligomerization sensor in the ADEP-bound MtClpP1P2 complex. The side chains of residues involved in salt-bridge formation are indicated. (E) Activity response curves, 40 °C, measured as a function of Bz-LL concentration using 250 µM PKM-AMC as substrate in the absence (blue curve) and presence (orange curve) of 25 µM ADEP.

### An allosteric framework for understanding MtClpP1P2 activation and function

The results of the previous section show that binding of ADEP to MtClpP2 leads to a shift in the activity profile of MtClpP1 that is most readily understood in the framework of an allosteric system. We next sought to address whether the binding of Bz-LL to the MtClpP1P2 complex was similarly allosteric. Initial NMR titrations made use of samples prepared with ILVM-labeled MtClpP1 (50 μM in protomer concentration) and a two-fold excess of [U-^2^H] MtClpP2, ensuring that more than 90% of the NMR-visible MtClpP1 ring forms a complex. Addition of Bz-LL leads to residue-specific titration profiles in which peaks present in the absence of ligand decrease in intensity concomitant with the appearance of new resonances, as would be expected for a system in slow exchange on the NMR chemical shift timescale (Fig. 4A – top three panels). Notably, nearly all of the ILVM methyl probes within the MtClpP1 ring respond to the addition of Bz-LL in this manner. A similar situation is observed for resonances derived from MtClpP2 when samples are prepared by reversing the labeling strategy indicated above (Fig. 4A – bottom three panels), and the titration profiles for MtClpP1 and MtClpP2 are essentially superimposable (see below). Because the spectrum of MtClpP2 is of relatively poor quality we did not attempt to assign individual correlations to specific sites in the protein, so that assignments for the selected residues in this ring are not available. Notably, the observed chemical shift changes were confined to the MtClpP1P2 complex; no changes to spectra of either isolated MtClpP1 or MtClp2 were observed when comparable amounts of Bz-LL were added, suggesting that the spectral perturbations report on a conformational change that only occurs in the full complex.

**Figure 4.**
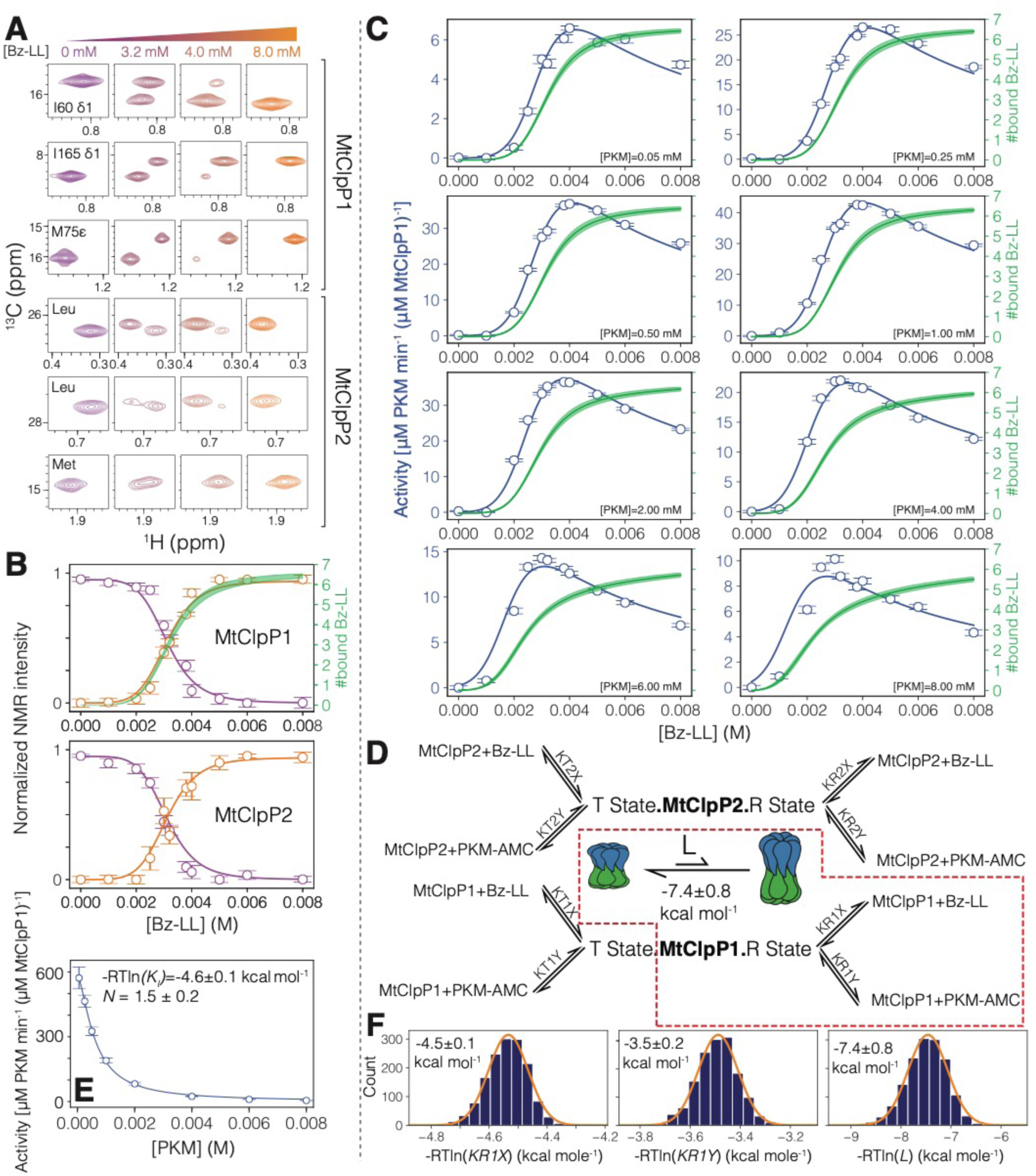
An allosteric network regulates the activation of MtClpP1P2. (A) Changes in correlations in ^1^H-^13^C HMQC spectra of MtClpP1P2 recorded as a function of Bz-LL concentration, 40 °C, 18.8 T, reflect interconversion between conformers on a slow timescale. The first three rows show spectral regions of datasets recorded on a MtClpP1P2 complex where MtClpP1 is ILVM-labeled (50 µM), while MtClpP2 is [U-^2^H] (100 µM). Assignments for MtClpP1 residues are indicated. The labeling pattern and protein concentrations are reversed in spectra of the bottom three rows. Assignments for MtClpP2 are not available and therefore only residue types are noted. (B) Changes in the intensities of *T* state (purple circles) and *R* state (orange circles) peaks averaged for resonances from MtClpP1 and MtClpP2 of the MtClpP1P2 complex. Error bars correspond to one standard deviation based on all peaks averaged. Solid lines are fits to a simplified version of the modified MWC model described in SI, whereby only the region highlighted by the dashed red box in panel D is included in the analysis. The number of Bz-LL molecules bound to MtClpP1 rings calculated from fits are shown (green line – thickness indicates an error of two standard-deviations from the mean; see SI). (C) Activity response curves (blue) measured at various Bz-LL and PKM-AMC concentrations. In each panel curves showing the number of Bz-LL molecules bound to the *R* state of MtClpP1 (solid green ± 2 standard deviations from the mean), calculated from the fits, are displayed. (D) Schematic representation of the derived MWC model, along with the simplified model used for global fitting of the NMR and functional data in this figure (red dashed box; See Fig. S8 for fitting of the complete model), including the *T*⇌*R* equilibrium, with equilibrium constant 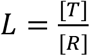. The relevant association constant are indicated in grey. (E) The decrease in *k*_*cat*_ as a function of substrate PKM-AMC concentration (open circles) fitted to a Hill-type substrate inhibition model (solid line). The fitted association constant (-RTln*K*_*i*_) for substrate binding to a distinct inhibitory site and the Hill constant (*N*) are listed. Errors are based on a Monte Carlo analysis discussed in SI. (F) The fitted free energies for Bz-LL and PKM-AMC binding to the *R* state of the MtClpP1 ring are listed, 40 °C, along with the distribution of values based on a Monte Carlo analysis. The distributions in panel (F) are fitted to a Gaussian function and the errors reported as twice the standard deviation in the parameters.

The decrease in the intensities of apo peaks (purple correlations in Fig. 4A) along with the concomitant increase in the intensities of the new set of correlations (orange) is quantified in Figure 4B for resonances from both MtClpP1 and MtClpP2 of the MtClpP1P2 complex. Here we have averaged the corresponding curves from each methyl probe (52 and 51 peaks from MtClpP1 and MtClpP2, respectively). When the averaged MtClpP1 and MtClpP2 profiles are fit separately to the simple Hill model for cooperative binding (solid lines), effective dissociation constants of 3.3 ± 0.2 mM (MtClpP1) and 3.1 ± 0.2 mM (MtClpP2) are obtained, along with Hill coefficients of 7.4 ± 0.9 and 7.9 ± 0.7 in each case, indicative of a highly cooperative transition.

The mM *K*_*d*_ values that have been fitted here are typically associated with binding events where the exchange kinetics are fast on the NMR chemical shift timescale. In this limit peak positions are shifted in a manner that reflects the relative populations of apo- and bound-conformers, without an increase in the number of peaks during the course of the titration that was observed here. In the present case an upper bound of 1-2 s^-1^ can be placed on the exchange rate between the observed pair of conformers based on the fact that exchange cross-peaks were not obtained in magnetization exchange experiments (61). Our results suggest, therefore, that the observed chemical shift behaviour does not correspond to a simple binding process. Indeed, separate titration profiles are obtained for a number of peaks that do shift in position as a function of [Bz-LL], as would be observed for a fast exchange, low affinity case. Unfortunately, we are not able to quantify these shifts properly to obtain a measure for the Bz-LL affinity because the shifts do not reach a plateau value at the concentrations used, and the changes in shift are small.

The slow-exchange process that is seen upon addition of Bz-LL (Figs 4A&B) most likely reflects a conformational change in the MtClpP1P2 complex that ensues upon Bz-LL binding, likely from the inactive, compact state that is observed in our cryo-EM derived structures (Fig. 2A&C) to the active, extended form that has been characterized by crystallography (53) (Fig. 2B&D). Functional assays performed over a range of PKM-AMC substrate concentrations are consistent with this interpretation (Fig. 4C – open circles). In these assays, MtClpP1P2 is inactive in the absence of Bz-LL, as expected since the handle region is not properly formed without activator (Fig. 2C). At ∼ 2.5 mM Bz-LL, MtClpP1P2 activity begins to increase sigmoidally until it reaches a maximum at ∼ 4 mM Bz-LL, followed by inhibition at higher Bz-LL concentrations (Fig. 4C). Notably, the rise in activity (Fig. 4C – blue circles) parallels the transition observed for both MtClpP1 and MtClpP2 as reported by the changes in peak intensities in NMR spectra (Fig. 4B – orange circles). There is thus a strong link between the NMR observations (Fig. 4B), the functional assays (Fig. 4C) and the structural changes characterized by cryo-EM (Fig. 2C).

To gain a quantitative understanding of MtClpP1P2 activation by Bz-LL we sought to simultaneously fit the NMR titration data (Fig. 4B – open circles) and the functional assays (Fig. 4C – open circles) to an allosteric binding model. One of the simplest and most intuitive models is that due to Monod, Wyman, and Changeux, termed the MWC model (62), and we have used a modified version that incorporates specific features of the MtClpP1P2 complex (Fig. 4D – see SI for derivation of the model). As in the original derivation we assume that each of the protomers of a given type, such as those from MtClpP1, for example, are equivalent, and that binding of substrate or activator at one site does not change the microscopic constants for binding at other locations. In addition, as described previously (62), all protomers can exist in one of two states, termed *T* (low activity) or *R* (high activity), with all seven protomers in a given ClpP ring in the same *R* or *T* conformation. A number of further assumptions is necessary: (i) As our Bz-LL titration data clearly establish that a conformational transition occurs simultaneously for both MtClpP1 and MtClpP2 (Fig. 4B, compare top and bottom), it is assumed that all protomers in both ClpP rings are in same conformation, either *T* or *R*. MtClpP1P2 tetradecamers where one of the ClpP rings is entirely *R* and the second entirely *T*, are, therefore, not allowed. (ii) Activity assays where the MtClpP1 ring is inactivated by mutation of the catalytic Ser in each of its seven protomers, MtClpP1(S98A)P2, show that maximum PKM-AMC hydrolysis rates are over ten-fold lower than for WT MtClpP1P2 (Fig. S7). We assume, therefore, that enzyme activity is solely governed by the number of substrate molecules bound to MtClpP1 that, in turn, is only active (capable of proteolysis) in the *R* state. (iii) Sites in MtClpP1 or MtClpP2 bind substrate and activator in a competitive manner, so that only one molecule of either substrate or activator is bound to a given site at a time. With these assumptions a set of equations can be derived that includes eight microscopic binding constants for the association of either substrate (PKM-AMC) or activator (Bz-LL) to protomers from either MtClpP1 or MtClpP2 rings, that in turn, are either both in *R* or both in *T* states (for example, *K*_*R1X*_ is the microscopic association constant for binding of Bz-LL to each of the seven protomers of the MtClpP1 ring of a MtClpP1P2 complex that is in the *R* state, Fig. 4D). In addition, an equilibrium constant 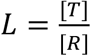, is required in the fits, where [*T*] and [*R*] are the concentrations of MtClpP1P2 in the *T* and *R* states in the absence of Bz-LL, respectively, as well as a *k*_*cat*_ value that ‘converts’ the average number of substrate molecules bound to the *R* state of the MtClpP1 ring to the measured substrate hydrolysis rates.

Global fitting of the NMR titration data (four curves; two for each of MtClpP1 and MtClpP2 rings – Fig. 4B) together with the functional assays at eight PKM-AMC and eleven Bz-LL concentrations (Fig. 4C) to a set of equations presented in SI produced poor fits of the data, especially for higher concentrations of PKM-AMC, where rates were predicted to be significantly larger than those observed. Next, we attempted to fit all of the activity profiles with a shared set of ligand association constants, as before, but using distinct *k*_*cat*_ values for each substrate concentration. Notably, in this case excellent fits were obtained, and interestingly the *k*_*cat*_ values so generated were reasonably well fit to a Hill curve, 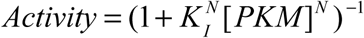, that takes into account the decrease in activity with substrate concentration (substrate inhibition, see below). Values of *N* = 1.5 ± 0.2 and *K*_*I*_ = 1.6 ± 0.3 × 10^3^ M^-1^ were obtained, where *K*_*I*_ is the association constant for the binding of one molecule of PKM-AMC to a distinct inhibitory site in each protomer, Figure 4E. Although this model yielded excellent fits of the experimental data over the complete range of substrate and activator concentrations used (Fig. S8), the majority of the parameters could not be obtained in a robust manner due to significant cross-talk between them, leading to large uncertainties in fitted values. We therefore sought a simpler version of the more general scheme of Figure 4D that would capture the behaviour of the system, with a smaller number of fitted parameters that could be extracted more robustly from the data. The simplest scheme that could fit the data is highlighted by the red box in Figure 4D, where only a single ring is considered corresponding to MtClpP1. It might be expected that a single MtClpP1 ring would be a good starting point for describing the activity profile since MtClpP1 is much more active than MtClpP2 (see above). Fits of the data to this simple model are indicated by the solid lines in Figures 4B&C. With the exception of the activity profiles at the highest substrate values ([PKM-AMC] = 6 mM and 8 mM) and for very low [Bz-LL] in these cases, remarkably good fits are obtained considering that only two association constants, corresponding to Bz-LL and substrate binding to the R state of MtClpP1, are used. Monte Carlo simulations were used to estimate the uncertainties in the extracted parameters and the distributions thus obtained for the free-energies of substrate and Bz-LL binding and for the *T*⇌*R* equilibrium are indicated in Figure 4F for the simple model used. As mentioned above, the quality of the fits can be further improved by introducing binding to the second ring. We have found that a more complex model allowing for substrate and activator binding to the *R* states of MtClpP1 and MtClpP2, but where binding does not occur to the *T* states of the rings (*i.e*., only the right half of the binding model of Fig. 4D is operational) improves the quality of the activity fits for the high substrate concentrations used.

A number of important conclusions emerge from the fits that are not dependent on the chosen model. First, the data can only be fit with large *L* values (∼2×10^5^), reflecting that in the absence of Bz-LL the compact and functionally inactive *T* state form of MtClpP1P2 is highly populated (∼99.999% for the case of *L* = 2×10^5^). Second, binding of Bz-LL shifts the *T*⇌*R* equilibrium to the active, extended *R* state, implying a higher Bz-LL affinity for the *R* state, and resulting in the observed sigmoidal NMR intensity and activity profiles. Third, at high Bz-LL concentrations the activity decreases, despite the increase in the population of *R*-state conformers, due to competitive inhibition between Bz-LL and substrate. The green profiles in Figures 4B and 4C show the average total number of bound Bz-LL molecules in the *R* state of MtClpP1 during the Bz-LL titration either without (Fig. 4B) or as a function of different concentrations of PKM-AMC (Fig. 4C), again assuming the simplest ‘single’ ring binding model. Fourth, given that the MtClpP2 ring has little activity towards the PKM-AMC substrate (Fig. S7), and that at maximum activity an MtClpP1 ring is bound with ∼ 4 Bz-LL molecules (Fig. 4C green line), most of the measured activity must derive from two or three unoccupied *R* state MtClpP1 protomers. Finally, the significant decrease in enzyme activity at higher substrate concentrations provides strong evidence for substrate inhibition in this system (see below).

### Allosteric activation is both intra- and inter-ring

As described above and illustrated in Figure 3E, the activity-response profile for MtClpP1P2 both increases and shifts to lower Bz-LL concentrations upon binding of ADEP. As ADEP only binds to MtClpP2 (48, 49) (Fig. 3A) while the measured activity derives from MtClpP1, these assays suggest that the adjacent rings of a complex are able to communicate with each other. There is strong evidence that allostery between rings extends to individual subunits within a ring as well. For example, fitted parameters from MWC models that consider binding of Bz-LL to the *R* states of both MtClpP1 and MtClpP2 rings of the complex establish that Bz-LL binds with at least ten-fold higher affinity to MtClpP1. Activation of MtClpP1P2 originates, therefore, predominantly from Bz-LL binding to the MtClpP1 ring with propagation from non-functional Bz-LL containing protomers to functional empty neighbouring subunits within the active MtClpP1 ring. To provide further evidence of intra-ring communication we have generated covalent complexes with the GLF-CMK mimic of substrate peptides that functions as an active-site inhibitor because of its covalent attachment to the MtClpP1 ring of MtClpP1P2. Intact protein electrospray ionization (ESI) mass spectra confirm that this inhibitor selectively modifies the MtClpP1 ring of the complex (Fig. S9), with no reactivity towards MtClpP2. As a first step, the structure of the MtClpP1P2 complex with the MtClpP1 ring fully reacted with GLF-CMK was determined using cryo-EM (Fig. S3E). After 2D and 3D classification (Fig. S3F), 143748 particle images were used in a final refinement with C7 symmetry enforced, producing a 3D map at a resolution of 3.5 Å (Fig. S4 E&F). An atomic model built into the cryo-EM map shows that MtClpP1P2 is in the extended conformation with a barrel height of 87 Å (Fig. 5A). Clear density for the GLF-CMK active-site inhibitor in each of the seven active sites of the MtClpP1 ring (Fig. 5B) is consistent with intact protein mass spectrometry (Fig. S9). Similar to the Bz-LL bound conformation, the handle region has a well-defined structure (Fig. 5C), and the oligomerization switch is engaged (Fig. 5D). However, unlike the Bz-LL bound form, density is not observed for the N-terminal domains of the protomers of the MtClpP2 ring in this fully modified MtClpP1P2 complex, suggesting that they are dynamic (Fig. 5B, Fig. S10).

**Figure 5.**
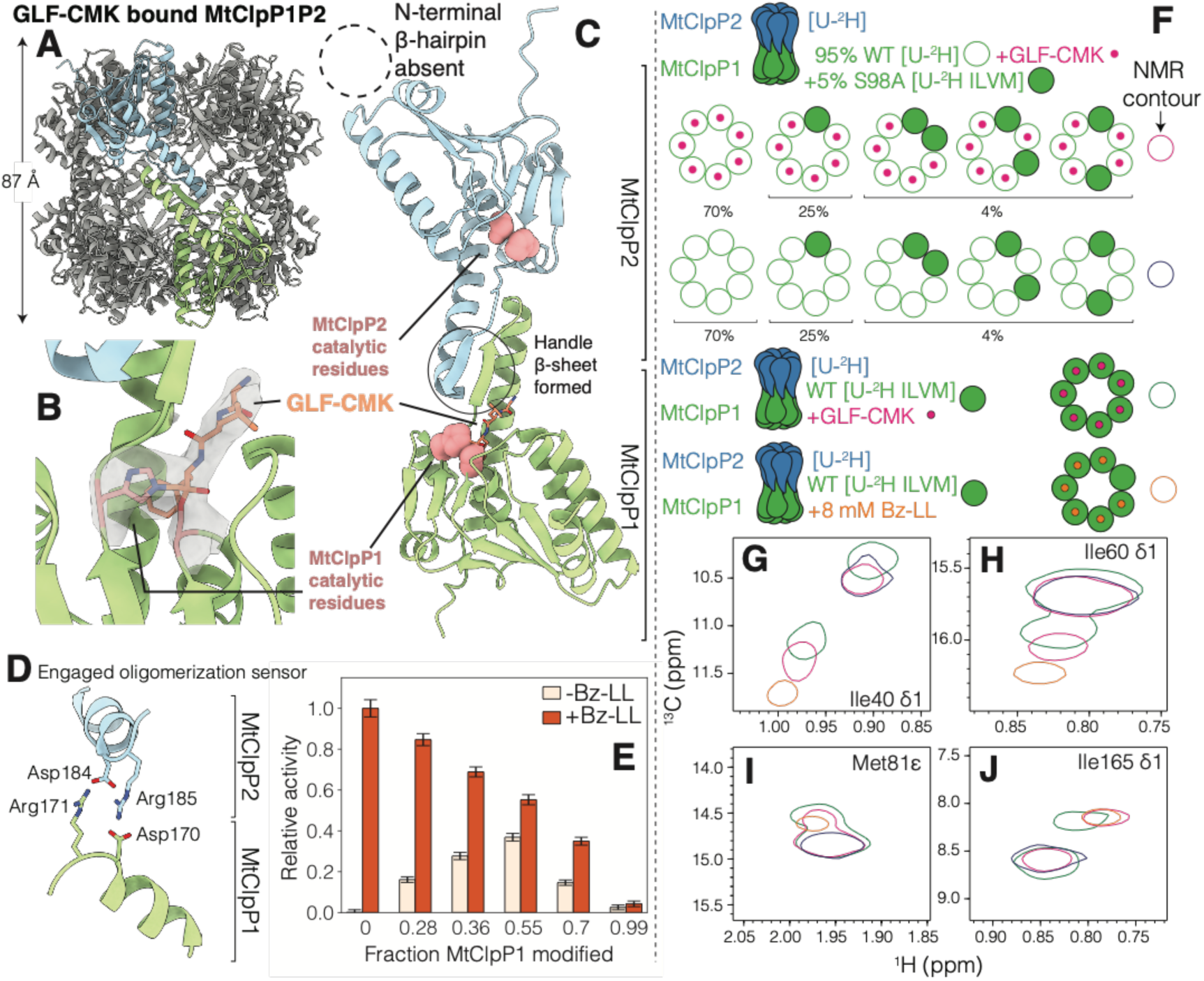
Active-site inhibitors activate MtClpP1P2 via an intra-ring allosteric communication. (A) Side view of the cryo-EM structure of MtClpP1P2 where the active sites of MtClpP1 are fully modified by GLF-CMK. The height of the MtClpP1P2 complex (excluding the N-terminal β-hairpins of MtClpP2) is noted. (B) Close up view of the catalytic His and Ser of an MtClpP1 protomer and cryo-EM density for the attached GLF-CMK (both His 123 and Ser 98 of the active site are covalently linked to the inhibitor). (C) Close-up view of a pair of opposite MtClpP1 (green) and MtClpP2 (blue) protomers showing an ordered handle β-sheet (solid circle) as a result of the modification of MtClpP1 by GLF-CMK. Notably, the N-terminal region of MtClpP2 (dotted circle) remains disordered. (D) Status of the oligomerization sensor of MtClpP1P2 in the GLF-CMK modified form. Side chains of residues involved in salt-bridge formation are indicated. (E) Activity of MtClpP1P2 complexes unmodified and partially modified with GLF-CMK assayed with 250 µM PKM-AMC substrate in the absence (light pink) and presence (orange) of Bz-LL (40 °C). The extent of the modification of the MtClpP1 ring of the MtClpP1P2 complex by GLF-CMK, measured from intact protein mass spectrometry (Fig. S9), is noted on the horizontal axis. Error bars correspond to one standard deviation from the average of three replicates. (F) (top row) Schematic of MtClpP1P2 complexes with hybrid MtClpP1 rings produced by mixing 95% [U-^2^H] WT (open green circles) and 5% ILVM-labeled S98A protomers (filled green circles), and their subsequent reaction with the GLF-CMK active-site inhibitor (filled pink circles). (second row) As above but without reaction with GLF-CMK. (bottom row) All MtClpP1 rings are either GLF-CMK reacted or partially Bz-LL bound (8 mM; orange circles. Note that approximately 6 Bz-LL molecules are bound to MtClpP1 at 8 mM, as indicated). The composition and population of the hybrid complexes (top two rows), predicted using combinatorial statistics, and the color schematic for panels G-J are displayed. (G-J) Isolated regions of ^1^H-^13^C HMQC spectra (40 °C, 18.8 T) of selected residues from ILVM-labeled MtClpP1 (pure and hybrid) in complex with unmodified, WT ^2^H-labeled MtClpP2 are shown. Assignments are indicated for each panel. Spectra are displayed as single contours to avoid cluttering.

To evaluate how covalent modification of the MtClpP1 ring impacts the activity of the MtClpP1P2 complex, that will provide evidence of intra-ring allostery, a series of protein samples where the MtClpP1 ring was partially modified by GLF-CMK was prepared by varying the ClpP:GLF-CMK ratio. The fraction of active sites modified (FM), defined as *FM* = *A*_*Mod*_/(*A*_*Mod*_ + *A*_*Unmod*_) where *A*_*Mod*_ and *A*_*Unmod*_ are the areas under the modified and unmodified ESI-MS protein peaks, respectively, was used to quantify the progress of the reaction (Fig. S9). Samples, ranging from no to full MtClpP1 modification were assayed for activity in the absence (Fig. 5E-light pink bars) and in the presence of Bz-LL (Fig. 5E-orange bars) using PKM-AMC as substrate. In the absence of GLF-CMK modification and without the addition of Bz-LL the MtClpP1P2 complex is inactive, consistent with the assays in Fig. 1B. Notably, MtClpP1P2 is activated at a FM of 0.28 without the addition of Bz-LL, with maximum activity achieved at FM ∼ 0.50, corresponding to modification of 3.5 out of 7 MtClpP1 protomers, on average. Thus, covalent modification of several catalytic S98 residues in the MtClpP1 ring leads to allosteric activation of the remaining protomers and generates a catalytically active enzyme. Further modification of the MtClpP1 ring reduces activity, and as expected complete modification abolishes catalysis. Interestingly, the addition of Bz-LL to these mixtures consistently results in further activation. In the presence of Bz-LL, maximum activity is observed when the MtClpP1P2 complex is not modified and decreases at higher levels of MtClpP1 modification with the concomitant reduction in the number of active sites.

The presence of intra-ring allostery can be further demonstrated by NMR studies of hybrid MtClpP1 rings produced by mixing 95% [U-^2^H] WT and 5% ILVM-labeled S98A protomers (Fig. 5F). The choice of a 95%-5% ratio maximizes the probability of generating rings with NMR-active protomers surrounded by GLF-CMK modified protomers (see below) so that the extent of intra-protomer communication within a ring can be assessed conclusively. The mixed rings are generated by unfolding WT and S98A protomers separately in guanidine hydrochloride (GdnCl), followed by mixing of the two chain types, and then subsequent refolding to generate complexes whose composition and population can be predicted using combinatorial statistics, as shown in Fig. 5F and described previously (63, 64). These composite MtClpP1 heptamers were then mixed with a two-fold excess of WT [U-^2^H] MtClpP2 heptamers and the complex reacted with GLF-CMK to modify all of the functionally active NMR-invisible WT MtClpP1 subunits (Fig. 5F-top panel, red dots). In contrast, NMR-visible S98A protomers do not react with GLF-CMK because they lack the catalytic serine; these ILVM-protomers serve as NMR reporters of whether intra-ring (within a MtClpP1 ring) allostery is present. A second unreacted complex, prepared in an otherwise identical fashion to the sample described above, and a third complex that includes IVLM-labeled WT MtClpP1 and [U-^2^H] WT MtClpP2, with the MtClpP1 fully reacted with GLF-CMK, were generated (Fig. 5F-bottom two panels). Spectra of fully modified MtClpP1 within the MtClpP1P2 complex showed pairs of peaks for many residues (green single contours), with one set of peaks at positions similar to those for the unreacted WT molecule (blue contours) and a second set of correlations that overlap well with those from the Bz-LL bound form of WT MtClpP1P2 (orange contours) (Figs. G-J). A similar set of pairs of peaks was observed for the mixed sample (NMR-invisible MtClpP2 + MtClpP1 comprised of largely NMR-invisible GLF-CMK protomers doped with NMR-visible MtClpP1 protomers; red single contours). These pairs of peaks report on an allosteric mechanism by which GLF-CMK binding to the active site of a protomer is transmitted to a second unbound subunit of the ring, as the NMR active protomers are not GLF-CMK labeled. The NMR data (Fig. 5F-J) and the activity assays of Figure 5E can be explained in the context of our proposed MWC model in which the covalent attachment of GLF-CMK partially shifts the *T*⇌*R* equilibrium towards the *R* state, in a similar manner to Bz-LL binding, although in the latter case the population can be completely shifted to the *R* conformation (Fig. 3B).

## Discussion

*M. tuberculosis* takes advantage of a tightly regulated Clp protein degradation system that helps maintain cellular homeostasis and allows the pathogen to remain dormant in the host for decades (27, 28, 44). It comprises a protease component (ClpP) that collaborates with ATP-dependent unfoldases, ClpX or ClpC1, to recognize, unfold, translocate, and ultimately destroy substrate proteins in the cell. The Clp system is essential for *M. tuberculosis* growth and virulence (26, 27), and as such its disruption by small-molecule modulators with antibiotic activity has generated considerable attention (29–32, 65, 66). ClpPs from actinobacteria, such as MtClpP1P2 studied in this work, are unique in that they are catalytically dead on their own *in vitro* but can be activated by the presence of a number of dipeptides, such as Bz-LL used here (32, 36, 49, 53). The structural basis for their lack of activity and their activation mechanism have remained poorly characterized but require a detailed understanding for the rational design and improvement of pharmaceuticals, such as those used to target MtClpP1P2. Quantifying the structure-function-dynamics paradigm in the context of MtClpP1P2 from *M. tuberculosis* forms the basis of our present study.

An intriguing feature of ClpP proteases is their structural plasticity and allosteric properties. It has been shown that ClpP, while maintaining its tetradecameric architecture, can assume several conformations that mainly differ in the height of its barrel-like structure and in the conformation of the handle domain (11, 67). High-resolution ClpP structures from various species include an active extended conformation with a fully formed catalytic triad and handle domain (38, 68), as well as compressed/compact forms with kinked/truncated handle helices and unstructured handle β-sheets with misaligned active site residues (69–72). While it is widely accepted that the extended form is prevalent in solution and is the active state, it is yet to be established whether the compressed/compact forms can be populated under conditions that are distinct from those used for crystallization, where crystal contacts can affect the ClpP structure (11, 73).

The potential of exploiting the inherent dynamics and allostery of ClpP to modulate its function has been well recognized (66, 74–77). One target is the handle region that mediates the interactions between protomers on opposite ClpP rings and anchors the catalytic residues to achieve optimal peptide bond cleavage. NMR measurements on *E. coli* ClpP revealed that the handle region is dynamic, interconverting between conformations that potentially allow product extrusion (78). Notably, cross-linking of pairs of opposing handle helices disrupted ClpP function by interfering with product release from the degradation chamber (78). While active site inhibitors such as Bortezomib are successful in inhibiting MtClpP1P2 activity and consequently the growth of *M. tuberculosis* (36), they lack specificity and lead to unwanted modification of serine proteases and the host proteasome (35). Novel noncovalent inhibitors exploit the dynamic nature of the handle helix to actuate a conformational change in the handle domain that distorts the catalytic triad of *S. aureus* ClpP (19). An N-terminal allosteric switch has been discovered via the introduction of a single V7A mutation in only one of fourteen identical protomers of the *S. aureus* ClpP tetradecamer, that leads to the formation of an inactive non-canonical split-ring conformation (64). Interestingly, this conformational change could be reversed and activity partially restored by ADEP binding. More recently Cediranib, an anti-cancer drug, was shown to bind to a hydrophobic pocket at the interface of the head and handle domains of MtClpP1, leading to inhibition of the MtClpP1P2 complex (37).

Our study provides an important step towards the goal of manipulating MtClpP1P2 function by providing the structural basis for why this complex is catalytically dead in the absence of activator peptides and why, unlike for many other ClpPs, binding of ADEPs do not activate the protein. The 3.0 Å resolution structure from cryo-EM of the apo-form of MtClpP1P2 shows that the complex is in the compact conformation. The hallmarks of the compact form are truncated handle helices and disordered handle β-strands that lead to an impaired catalytic triad, thereby explaining the lack of activity observed for apo-state of the complex. Notably, while ADEP binding to MtClpP1P2 leads to an ordering of the N-terminal region of MtClpP2, the handle region remains disordered so that the active site residues are not properly positioned with respect to each other for catalysis. The binding of ADEP, as described here, as well as either ClpA or ClpC1 to MtClpP1P2 (28, 53), results in a significant shift in the MtClpP1P2 activity response curve to lower Bz-LL concentrations, suggesting that the concomitant use of active site inhibitors together with ADEP might increase the susceptibility of *M. tuberculosis* to antibiotic treatment.

The pervasive contributions of allostery in the MtClpP1P2 system is highlighted by our study. The binding of ADEP exclusively to sites in MtClpP2 leads to enhanced catalytic activity of active sites in MtClpP1 that is only possible via inter-ring communication. Addition of the active-site inhibitor GLF-CMK to generate a fraction of MtClpP1 protomers that are modified at catalytic serine residues resulted in activation of an otherwise catalytically dead enzyme, suggesting an important role for intra-protomer communication within a single ring. The high degree of allostery has been captured by a modified MWC model that reproduces how activator peptide Bz-LL binding to MtClpP1P2 leads to the conversion from a catalytically dead *T*-state to an active *R*-conformation, as observed through large chemical shift perturbations in NMR titration spectra, and accounts for the activity *vs* Bz-LL concentration profiles as a function of PKM-AMC substrate concentration. The functional assays show a marked decrease in activity with increasing substrate concentration past a certain threshold that can be accounted for by invoking substrate inhibition and taken into account in our MWC model (see SI). A model of substrate rather than product inhibition is more likely for a number of reasons. First, substrate degradation kinetics were linear over the complete time course examined, while product inhibition would lead to decreased rates over time. Second, activity profiles decreased with increasing substrate concentration beyond PKM-AMC substrate concentrations of ∼ 1 mM, despite the fact that the amount of product produced also decreased. It is estimated that in ∼20% of enzymes the substrate may serve as an inhibitor (79), controlling enzyme activity during periods of unusually low or high metabolic activity. Substrate activation and inhibition might be advantageous for an organism like *M. tuberculosis* with a slow growth rate that must retain viability during latent infections (80). Downregulation of MtClpP1P2 activity in response to extreme substrate concentrations, either low or high, could reduce spurious degradation of small disordered proteins or newly-synthesized polypeptides which have yet to fold properly that diffuse into the degradation chamber independent of regulatory unfoldases.

Although our model accounts for all of the measured experimental data, at least when substrate and activator binding to both rings of the complex is considered (Fig. S8), we have not been able to cross-validate the presence of an independent site that binds substrate and leads to inhibition. Titration of substrate into a sample with active MtClpP1P2 results in rapid hydrolysis. Generation of a catalytically dead enzyme by substituting active-site Ser residues in MtClpP1 and MtClpP2 with Ala circumvents this problem, but generates enzymes with properties that are different from those of the WT. For example, the *T* to *R* transition occurs at Bz-LL concentrations that are 5-fold reduced for the Ser to Ala mutant, and a complex that is comprised of WT MtClpP1 and S110A MtClpP2 has an activity profile that is different from the WT enzyme with a shift in activity to lower Bz-LL concentrations.

Although binding of active-site inhibitors has been reported to activate *Listeria monocytogenes* ClpP (81) and the β-subunits of the proteasome with trypsin-like activities (82), a mechanistic understanding of this phenomenon has been lacking. Previous studies made use of a multi-pronged approach involving isothermal calorimetry, NMR spectroscopy, crystallography, and functional assays to understand the mechanism of allosteric activation of *Thermus thermophilus* ClpP (TtClpP) (83). It was shown that non-saturating concentrations of the active site inhibitor Bortezomib leads to an activation that is moderately cooperative, with an activity profile that could be fit with a Hill coefficient of 1.3. Notably, Bortezomib binding resulted in opening of the N-terminal axial pores that enabled the passage and degradation of larger substrates by TtClpP. The structural differences between the conformers involved in the allosteric transition could not be established experimentally, however, as both the apo and the Bortezomib-bound forms crystalized in the extended conformation. However, the authors used MD simulations to conclude that the less active form is likely the compressed state of the complex, with kinked handle helices.

The allosteric model of ClpP activation based on an equilibrium between active and inactive conformations is useful to understand why some ClpPs, such as those from *T. thermophilus* and *M. tuberculosis*, exhibit non-Michaelis Menten kinetics and cooperative activation and why others such as *E. coli* ClpP have unusually high substrate proteolysis rates with no requirement for activator peptides (36). In the framework of the MWC model the equilibrium between *T* and *R* states would be shifted much more closely to *R* for ClpP from *E. coli* than for TtClpP that in turn is significantly closer to *R* than is the case for MtClpP1P2. Our modeling of Bz-LL binding data and activity profiles suggest that 0.0005% of the MtClpP1P2 molecules in solution are in a catalytically competent state in the absence of activator. By contrast, approximately 25% of TtClpP molecules are in an active state. The extent of allostery likely represents yet another layer of regulation of ClpP activity that is carefully tuned in each organism. In the case of dormant bacterial populations of *M. tuberculosis*, cellular homeostasis would require a tight regulation of all components of the protein degradation machinery so that spurious substrate removal is limited (13, 14). This regulation is partly achieved through the binding of ClpC1 or ClpX regulatory particles, which is coupled to ClpP protease active sites (28, 44). A second layer of protection against unwanted degradation is afforded by the modest affinities of MtClpP1 and MtClpP2 for each other, and the fact that individual ClpP rings are catalytically dead since interactions between Asp and Arg residues on adjacent rings are required to couple the oligomerization state of the enzyme to its catalytic activity. Finally, a third layer of control is provided by allostery, allowing MtClpP1P2 to adjust its activity in response to high concentrations of substrates. In this way the enzyme remains inactive, even in a tetradecameric form, unless a certain activating threshold of substrate is achieved. For Bz-LL and GLF-CMK it appears that, on average, binding of roughly four molecules is needed to trigger this process. Although high concentrations of Bz-LL activator are required to convert to the catalytic state it is worth noting that the presence of a single translocated substrate trapped in the catalytic chamber of MtClpP1P2 with an internal volume of 65600 Å^3^ (84) corresponds to an effective concentration of ∼ 25 mM that could be sufficient to promote activity. Thus, the activation mechanism described here may also occur *in vivo* where both the binding of AAA+ unfoldases and the translocation of substrates bearing activating sequences lead to the stabilization of the extended form of MtClpP1P2 so that efficient and processive substrate degradation is enabled.

## Materials and Methods

Details of protein expression and purification, biochemical and biophysical measurements, along with data fitting and derivation of analytical models are provided in Supporting Information.

## Acknowledgements

We thank Dr. Samir Benlekbir for assistance with cryo-EM data collection on the Titan Krios, Professor Walid A. Houry (University of Toronto) for the gift of ADEP, Professor Eric J. Rubin, Drs. Tatos Akopian, and Olga Kandror (Harvard) for an initial supply of Bz-LL and valuable discussions, Drs. Algirdas Velyvis, Jacob P. Brady, and Justin M. Di Trani for help with data fitting, and Dr. Robert Vernon for valuable discussions and assistance with Rosetta modelling. ZAR and SV were supported by a scholarship and a postdoctoral fellowship from the Canadian Institutes of Health Research, respectively. LEK and JLR were supported by the Canada Research Chairs program. This research was funded by Canadian Institutes of Health Research grants MOP-133408 (LEK) and PJT-162186 (JLR). Titan Krios cryo-EM data were collected at the Toronto High-Resolution High-Throughput cryo-EM facility supported by the Canadian Foundation for Innovation and Ontario Research Fund.

## SI Appendix

### Materials and Methods

#### Plasmids and constructs

The *clpP1* (Uniprot: P9WPC5) and *clpP2* (Uniprot: P9WPC3) genes from *Mycobacterium tuberculosis* and the *clpP* gene from *S. aureus* (Uniprot: A6QF76) were synthesized by GenScript (Piscataway, NJ, USA) and cloned into the NdeI and BamHI sites of pET24a+ (Novagen, Madison, WI, USA). A cleavable N-terminal His_6_-SUMO tag was introduced into all constructs via Gibson assembly. Point mutations were introduced using Quikchange site-directed mutagenesis (Agilent, Santa Clara, CA, USA).

#### Protein expression and purification

MtClpP1, MtClpP2, and SaClpP were expressed heterologously and purified as detailed in previous work (1). Briefly, transformed BL21(DE3) Δ*clpP∷cat E. coli* cells were grown in minimal M9 D_2_O media supplemented with ^15^NH_4_Cl and d_7_-glucose as the sole nitrogen and carbon sources, respectively. Cells were grown at 37 °C and protein overexpression was induced with 0.1 mM IPTG at OD_600_=1.0 and allowed to proceed for ∼18 hours at 18 °C. [U-^2^H; Ileδ1-^13^CH_3_; Leu,Val-^13^CH_3_/^12^CD_3_; Met-^13^CH_3_]-labeled (referred to a ^13^CH_3_ ILVM-labeled in text) and Val/Leu-γ1/δ1(proR) samples were produced as described previously (1). For the production of samples for cryo-EM analysis, cells were grown in Lysogeny Broth (LB) media at 37 °C and induced at OD_600_=1 with 0.1 mM IPTG. Expression was allowed to proceed for 18 hours at 25 °C. Proteins were first purified using Ni-affinity chromatography. The Ni lysis/wash buffer contained 50 mM Tris, 300 mM KCl, 20 mM imidazole, and 10% glycerol adjusted to pH 7.0. The imidazole concentration was increased to 500 mM in the Ni elution buffer. Because MtClpP2 tends to aggregate at high imidazole concentrations, elution from the NiNTA column was collected in a Falcon tube containing ∼25 mL of Ni lysis/wash buffer to immediately dilute the high imidazole concentration in the Ni elution buffer. This step was followed by cleavage of the SUMO tag using Ulp1 protease. This mixture was concentrated using an Amicon Ultra-15 50K MWCO concentrator and subjected to size exclusion chromatography (SEC) on a Superdex 200 Increase 10/300 (GE Healthcare) column in SEC Buffer containing 50 mM imidazole, 100 mM KCl, 5 mM DTT adjusted to pH 7.0. Protein concentrations were determined spectrophotometrically (GdnCl-denatured protein) using extinction coefficients obtained from ExPASy’s ProtParam web-based tool (https://web.expasy.org/protparam/). For NMR measurements and degradation assays, the samples were buffer exchanged into NMR Buffer containing 50 mM imidazole, 100 mM KCl, 1 mM TCEP adjusted to pH^measured^ 7.0 prepared in 99.9% D_2_O.

#### Preparation of mixed MtClpP1P2 complexes for NMR

MtClpP1 rings containing mixtures of WT and S98A protomers were prepared following a procedure described previously (1, 2). Briefly, pure [U-^2^H] WT and ILVM-labeled S98A MtClpP1 heptamers were purified separately and mixed to achieve a 95%:5% WT:S98A ratio. This protein mixture was concentrated to ∼0.5 mL and unfolded and diluted with the addition of unfolding buffer containing 100 mM KCl, 50 mM imidazole, 6 M GdnCl, and 10 mM DTT at pH 7.0 to a final protein concentration of 500 μM. The complexes were reconstituted by drop-wise addition into a refolding buffer containing 100 mM KCl, 50 mM imidazole, 1 M arginine, 10 mM DTT, and 15% glycerol at pH 7.0 to a final GdnCl concentration of 300 μM. The refolded mixture was concentrated to ∼1 mL using an Amicon Ultra-15 50K MWCO (Millipore) concentrator and then applied to a Superdex 200 Increase 10/300 (GE) column in SEC buffer. The resultant MtClpP1 rings eluted in a manner identical to that of pure WT MtClpP1 and there was no protein in the void volume. MtClpP1 fractions were pooled and mixed with a two-fold excess of [U-^2^H] WT MtClpP2, and allowed to reacted overnight with a ten-fold excess of benzyloxycarbonyl-Gly-Leu-Phe-chloromethyl ketone (abbreviated GLF-CMK throughout the manuscript) (New England Peptide Inc, Gardner, MA, USA) in the presence of 4 mM Bz-LL. Following verification of complete GLF-CMK modification of MtClpP1 active sites by ESI-MS, Bz-LL was removed by buffer exchanging the mixture into NMR Buffer.

#### NMR Spectroscopy

All NMR measurements were performed at 18.8 T and 40 °C, using a Bruker AVANCE III HD spectrometer equipped with a cryogenically cooled x,y,z pulsed-field gradient triple-resonance probe. ^1^H-^13^C correlation spectra were recorded as HMQC datasets, exploiting a methyl-TROSY effect that is particularly beneficial for applications to high molecular weight proteins (3). Spectra were processed using the *NMRPipe* suite of programs (4), analyzed using scripts written in-house, and visualized using *Ccpnmr* (5).

#### Methyl group assignment

Over 90% of the Ile (17/17), Leu (14/18), Met (8/8), and Val (7/7) correlations in spectra of MtClpP1, 40 °C, were assigned by a combined mutagenesis and NOE-based approach, as described previously (1, 6). ^1^H-^13^C HMQC spectra were recorded of the following MtClpP1 assignment mutants: M75L, M81L, M122L, I29V, I30V, I40V, I60V, I77V, I88V, I120V, I136V, I189V, V36I, V82I, V129I, V145I, V186I, V195I, L14M, L16M, L25M, L44M, L50M, L83M, L103M, and L126M. A methyl-TROSY based 3D NOE experiment (7) that records chemical shifts as ^13^C[i]-NOE-^13^C[j]-^1^H[j] was measured on an ILVM-labeled sample of apo WT MtClpP1 with an NOE mixing time of 250 ms. To extend assignments to the *T* and *R* states of MtClpP1P2, NOE experiments were also recorded on samples that contained 1 mM ILVM-labeled MtClpP1 mixed with 1.2 mM [U-^2^H] MtClpP2 in the presence or absence of 8 mM Bz-LL. The side chains of Leu in Bz-LL were uniformly deuterated to remove T1 noise. Stereospecific assignments of methyl groups of Leu and Val residues was obtained using the labeling approach described previously (8).

#### NMR data fitting

Binding constants for the MtClpP1 and MtClpP2 interaction (with and without 8 mM Bz-LL) were obtained by titrating [U-^2^H] MtClpP2 into an ILVM-labeled MtClpP1 sample (50 μM in monomer concentration). The decrease in the intensities of cross-peaks derived from the unbound state and the concomitant increase in cross-peak intensities from the bound-state can be quantified and subsequently fit as a function of MtClpP2 concentrations to the following expressions,

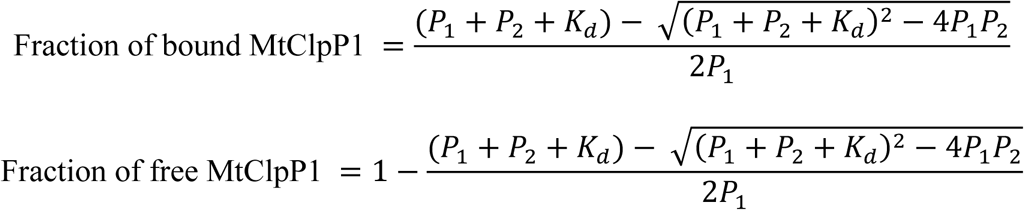

where P_1_ and P_2_ are the total concentrations of MtClpP1 and MtClpP2 at each titration step, respectively, and *K*_*d*_ is dissociation constant. Peak intensities were extracted using the *NMRPipe* suite of programs (4) and data-fitting was performed using a Python script written in-house.

#### An extended MWC model that includes competitive binding of Bz-LL and substrate

We begin by initially considering a binding model for a simplified system comprising a pair of rings, as for MtClpP1P2, but where each ring in turn contains of only a single binding site for activator (denoted as *X*) or substrate (*Y*). As described in the text, we assume that binding of *X* and *Y* is competitive (*i.e*., both bind to the same site) and that each ring is in the same state, either *R* or *T*. We distinguish rings 1 and 2 by subscripts 1, 2, so that *R*_*1*_*R*_*2*_ denotes a complex where rings 1 and 2 are both in the *R* state, for example. A number of equilibria immediately follow,

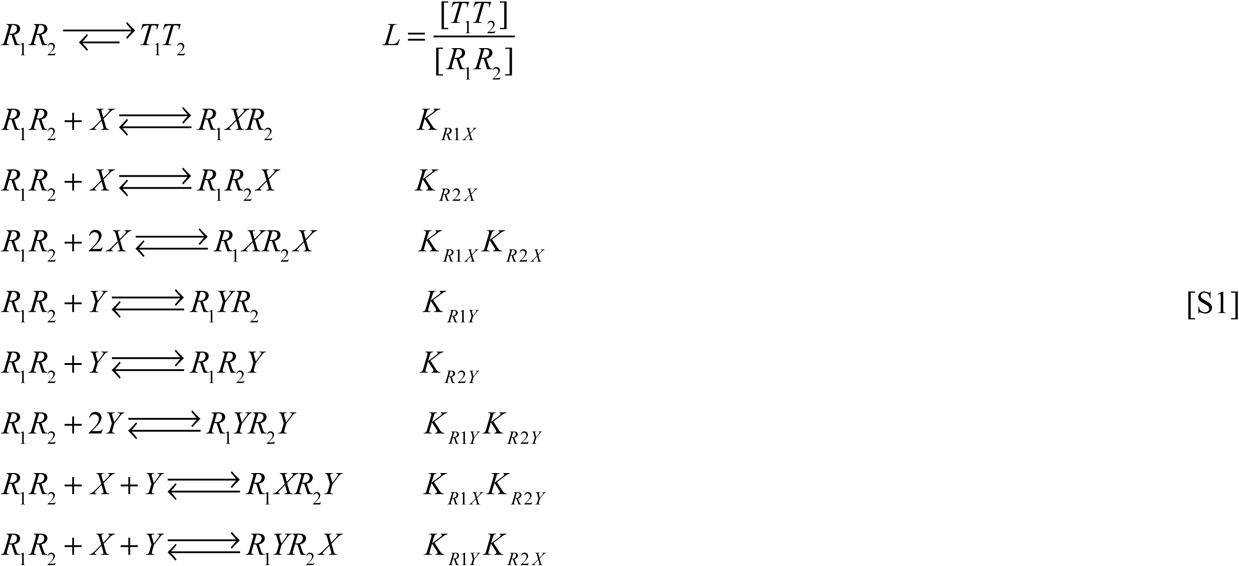

A similar set of equations exists for binding of *X* and *Y* to *T*_*1*_*T*_*2*_ that are obtained by replacing each *R* with *T* in Eq. [S1]. We now define a binding polynomial *Q* that is equal to the sum of the concentrations of all protein components,

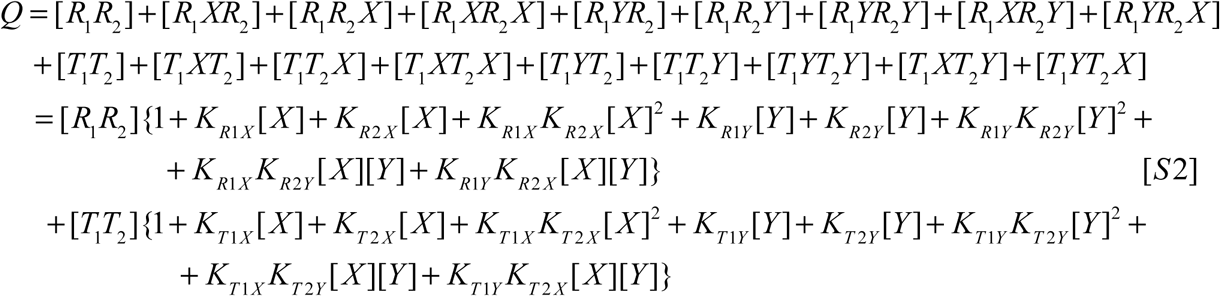

where *R*_*1*_*XR*_*2*_ is a complex with ligand *X* (Bz-LL) bound in ring 1, *R*_*1*_*XR*_*2*_*Y* is a complex with ligands *X* and Y (substrate) bound to rings 1 and 2, respectively, and so forth. Eq [S2] can be simplified using the first entry of Eq. [S1] to give,

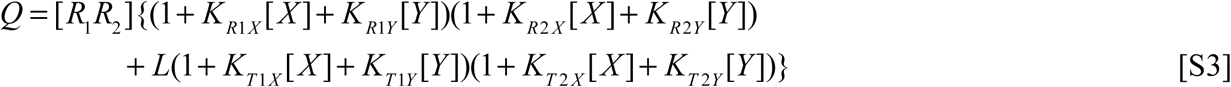

The fraction of conformers where both rings are in the R state, *f*_*R*_, is given by the sum of the first 9 terms of Eq. [S2], ([*R*_1_*R*_2_] + [*R*_1_ *XR*_2_] + … + [*R*_1_*YR*_2_ *X*]) divided by the total concentration of protein, *Q*,

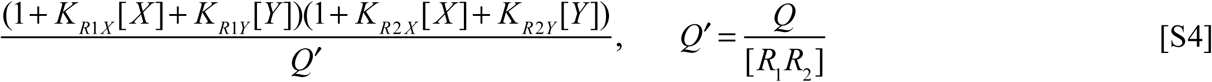

and similarly the fraction of conformers where both rings are in the T configuration, *f*_*T*_, is

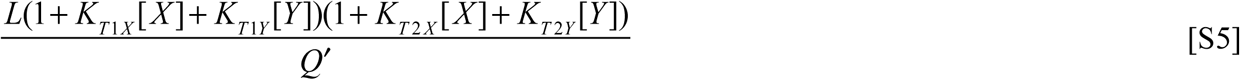

Assuming that the activity of the complex (*A*) is proportional to the average number of substrate molecules bound to *R*_*1*_, as discussed in the text, it follows that

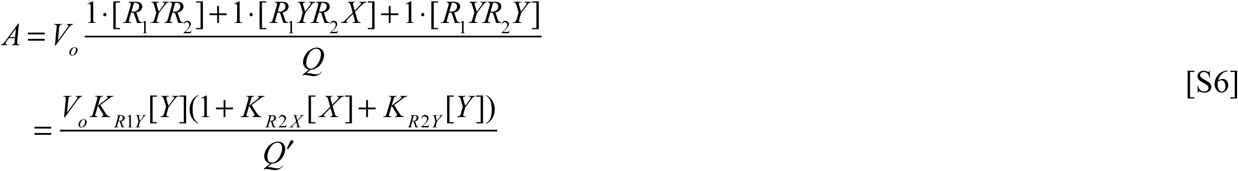

where the constant of proportionality *V*_*o*_ is the activity measured under saturating amounts of *Y* (limit when *K*_*R1Y*_[*Y*]>>1), when [*X*]=0 and assuming all molecules are in the active *R* state. The power of the binding polynomial approach (9) that has been adopted here is that expressions like Eq [S6] can be readily derived directly from *Q* (or *Q*′) Consider the case where *Q*′ is given as

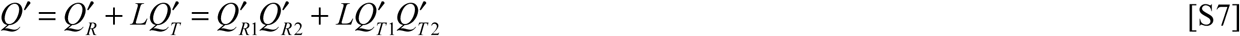

where 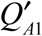 and 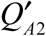 contain terms related only to the binding of *X* or *Y* to rings 1 and 2, respectively, and *A* ∈ {*R, T*}, corresponding to the *R* or *T* state of a ring. In the case of the example here

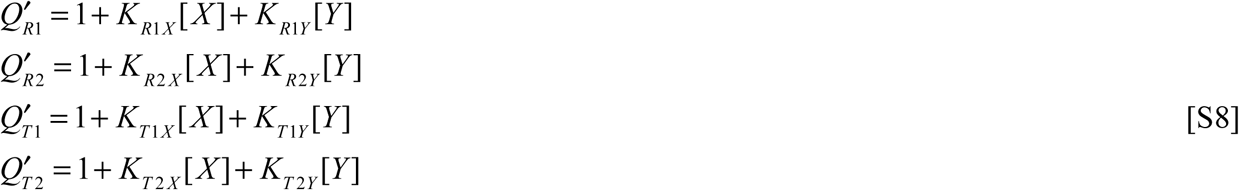

and the average number of substrate molecules bound to the *R* state of a complex, *R*_*1*_*R*_*2*_, (both rings are in the *R* state) is given by (9)

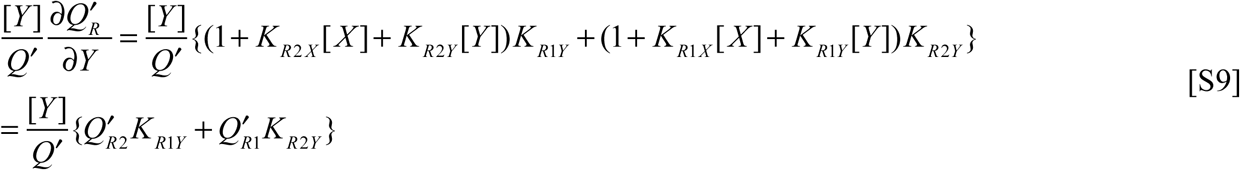

while the average number of substrate molecules bound to *R*_*1*_ can be derived from the relation 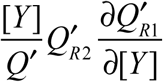 to give the first term of Eq. [S9] which is the expression in Eq. [S6], neglecting *V*_*o*_.

The results of this section can be easily generalized to the case where each ring is made up of *n* protomers (*n*=7 for MtClpP1P2), rather than 1 considered to this point. In this case,

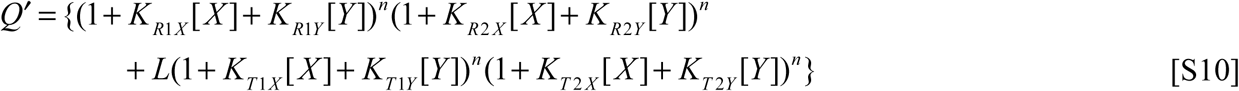

and *f*_*R*_ and *f*_*T*_ are given by

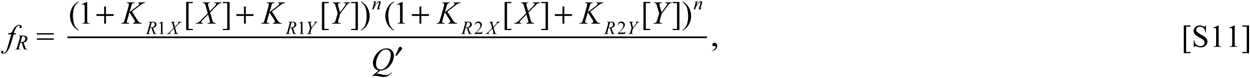

and

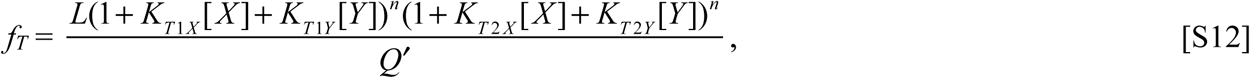

respectively. Similarly, *A* can be expressed as

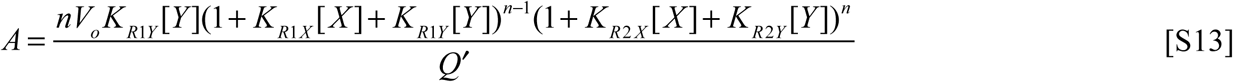

where *V*_*o*_ is the maximal activity per subunit. As described in the text, we were not able to fit our functional assays using Eq. [S13]. The data were well fit, however, when *V*_*o*_ was allowed to vary with [*Y*], that is *V*_*o*_ = *V*_*o*_([*Y*]), that takes into account the decrease in *A* at high concentrations of substrate (substrate inhibition). Thus, in the fits illustrated in Figure 5C *V*_*o*_([*Y*]) is floated for each [*Y*] (each panel). The plot of Figure 5E shows *V*_*o*_[*Y*] *vs* [*Y*] (circles) and a fit to the data using the function 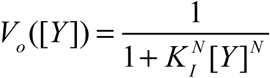 (solid line) where *K*_*I*_ is the association constant for the binding of each of *N* molecules of *Y* to inhibitory sites on ClpP1P2 and *N* is the Hill coefficient for this process,

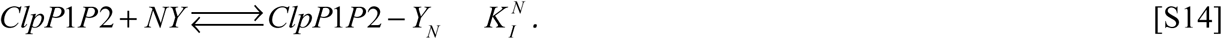

Eqs. [S11], [S12] and [S13] (where *V*_*o*_ = *V*_*o*_([*Y*]) have been used to obtain the fits shown in Figures 4B&C of the main text, with scaling factors included for *f*_*R*_ and *f*_*T*_ that are used to weight the NMR data relative to the data from the functional assays. In all fits, the four NMR intensity curves of Figure 4B have been multiplied by 14 prior to fitting, to account for the larger amount of activity data and activity values that can peak at approximately 30.

The average number of activator (Bz-LL = *X*) molecules that are bound to MtClpP1P2 can be calculated as follows,

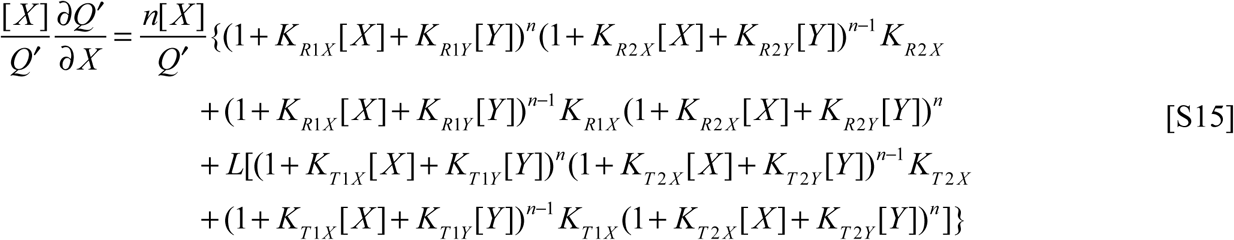

where *Q*′ is given by Eq [S10]. In a similar manner, the average number of activator (Bz-LL = *X*) molecules that are bound to the *R* configuration of the complex (i.e., both rings are in the *R* state) is calculated as,

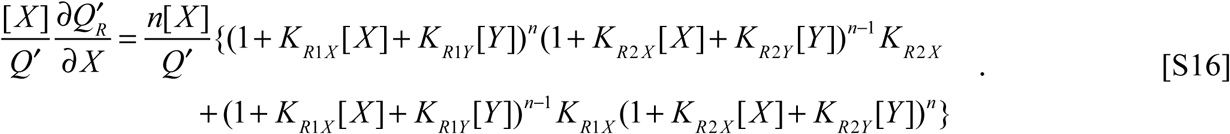

Finally, the average number of activator molecules that are bound to the MtClpP1 ring in the *R* configuration of the complex (green solid curves in Figures 4B&C) is

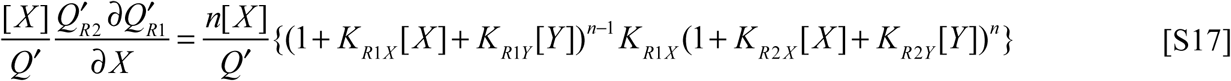

#### Activity Assays

##### Peptidase rate measurements

The peptidase activity of MtClpP1P2 was measured at 40 °C with Acetyl-L-Pro-L-Lys-L-Met bearing a C-terminal fluorogenic 7-amino-4-methylcoumarin group (abbreviated PKM-AMC throughout the manuscript) as substrate. The reaction was followed with a Synergy Neo2 96-well microplate reader by taking a measurement every 21 seconds for 60 minutes at λ_ex_: 355 nm, λ_em_: 460 nm. For all activity response measurements the concentration (protomer) of MtClpP1 was 1 µM while the concentration of MtClpP2 (protomer) was 20 µM, ensuring that the fraction of MtClpP1 in complex with MtClpP2 does not change substantially as [Bz-LL] is varied (note the difference in K_d_ values when Bz-LL is added, Fig. 1E&F). To ensure maximum similarity with the NMR titrations and facilitate subsequent data fitting, all functional assays, except for those shown in Figure 5, were performed using deuterated enzyme in 100% D_2_O-based NMR buffer. Activities are derived from initial rates extracted and analyzed using a Python script written in-house. Error bars correspond to one standard deviation derived from three repeat measurements.

#### Combined analysis of NMR intensities and activity assays as a function of Bz-LL concentration (Fig. 4B-F)

Peak intensities in ^1^H-^13^C HMQC spectra of MtClpP1P2, with ILVM-labelled ClpP1 or ClpP2, were quantified as a function of 11 Bz-LL concentrations (Fig. 4B) and activity profiles of MtClpP1P2 measured from initial rates of fluorescence change due to hydrolysis of the substrate PKM-AMC quantified with 11 Bz-LL concentrations and 8 substrate concentrations. The NMR and activity data were jointly fit to a modified MWC model (Fig. 4D, described in SI) using Eqs. [S11]]-[S13] with *V*_*o*_([*Y*]) floated for each concentration of *Y*. This was accomplished using a protocol in which initial guesses for parameters were derived by a grid search that explored values for the association constants between 10^−4^ M^-1^ and 10^4^ M^-1^, between 10^−2^ A.U. and 10^6^ A.U. for *V*_*o*_[(*Y*)] (A.U. = μM PKM min^-1^/μM MtClpP1P2) and between 10^−2^ and 10^6^ for *L*. Each dimension was searched in 8 steps that were linear on the logarithmic scale. Once starting parameters were identified by this procedure a least squares fit of the data was performed using an in-house package written in Python 3.8 using a Levenberg-Marquardt search engine that is available within the lmfit (v. 0.9.14) package (10). The 8 extracted values of *V*_*o*_([*Y*]) were subsequently fit to a Hill-model of substrate inhibition described above to obtain the association constant for substrate binding to the inhibitory sites in the complex. Errors in the fitted parameters were estimated using a Monte Carlo approach whereby random errors, calculated as the root-mean-squared deviation between experimental points and those generated with the best-fit model, were added to the best-fit model to produce 1500 synthetic data sets which were fit as per the experimental data. Initial guesses for the Monte Carlo runs were generated randomly. The values derived from Monte Carlo repeats were converted to a histogram, which was subsequently fit to a normal distribution function to yield expectation values and standard deviations *σ*. Final errors are reported as 2*σ* in the extracted values (95% confidence interval).

#### Cryo-EM

##### Sample preparation for cryo-EM

All purified proteins were concentrated to ∼20-30 mg/mL in buffer. Immediately before freezing, samples were mixed with 0.025 % (wt/vol) IGEPAL CA-630 (Sigma-Aldrich) to increase the proportion of protein complexes adopting side views on the grid. 2.5 μL of the sample mixtures were applied to nanofabricated holey gold grids (11–13) with a hole size of ∼1-2 μm. Grids were blotted on both sides using a FEI Vitrobot mark III for 15 seconds at 4 °C and ∼100% relative humidity before freezing in a liquid ethane/propane mixture (14).

##### Electron microscopy

All MtClpP1P2 complexes were imaged with a Thermo Fisher Scientific Titan Krios G3 microscope operating at 300 kV and equipped with a FEI Falcon III DDD camera. Structures were calculated from counting mode movies consisting of 30 frames, obtained over a 60 second exposure with defocuses ranging from 0.7 to 2.0 μm. Movies were at a nominal magnification of 75000×, corresponding to a calibrated pixel size of 1.06 Å and with an exposure of 0.8 electrons/pixel/s, giving a total exposure of 43 electrons/Å^2^. For apo, ADEP-bound, and GLF-CMK modified MtClpP1P1 2092, 725, and 1645 movies were collected respectively, using the microscope’s *EPU* software.

##### EM image analysis

Whole frame alignment was performed in *cryoSPARC* v2 (15) with the resulting averages of frames used for contrast transfer function (CTF) determination (16). Templates for particle selection were generated by 2D classification of manually selected particles. Particle images were extracted in 184×184-pixel boxes, and individual particle alignment and exposure weighting was performed within *cryoSPARC* v2 (15).

##### Atomic model building and refinement

To model MtClpP1P2 (S10), a single subunit of each of MtClpP1 and MtClpP2 of the Bz-LL bound crystal structure (PDBID: 5DZK) (17) was fit into the EM density map as rigid bodies using *UCSF Chimera* (18). For apo MtClpP1P2, *Rosetta* (19) was used to minimize the structure with C7 symmetry enforced, with iterative backbone rebuilding. The best scoring models were visually inspected, and the best fitting model was used for further analysis. For ADEP bound MtClpP1P2 a single pair of protomers, one from MtClpP1 and one from MtClpP2 (apo structure) were used as a starting model for further refinement. The N-terminal domains were built in *Coot* (20), and the entire model relaxed with *Rosetta* enforcing C7 symmetry. The top scoring models were then used for further analysis. ADEP was modelled based on PDBID 6CFD, in *Coot*, and real space refined in *PHENIX* using ligand restraints built in *PHENIX* elbow (21). For GLF-CMK bound MtClpP1P2, MtClpP1 and MtClpP2 protomers from the Bz-LL bound crystal structure (PDBID: 5DZK) were rigidly docked into the density in *UCSF Chimera*. Restraints for the GLF-CMK ligand were generated in *PHENIX* elbow followed by real space refinement in *PHENIX*. Validation reports (Table 1) were prepared in *PHENIX.*

Models were evaluated with *Molprobity* (22) and *EMRinger* (23). Figures were generated in *UCSF Chimera* (18) and *UCSF ChimeraX* (24), and colors chosen with *ColorBrewer* (25).

---

**Figure S1.**
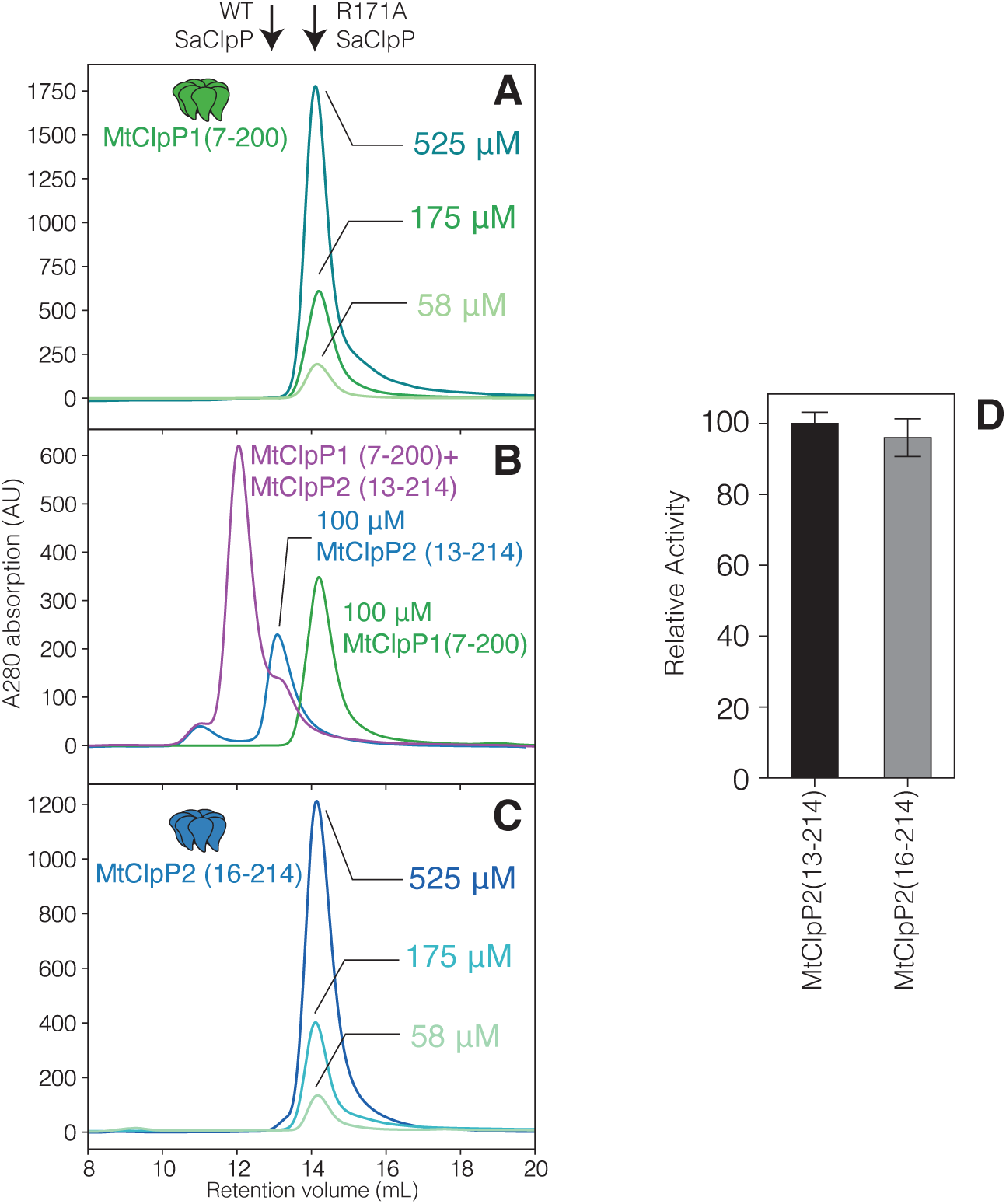
Effect of protein concentration on oligomeric state of (A) MtClpP1 (residues 7-200) as established by SEC. (B) SEC profile of MtClpP2 with native propeptide processing (residues 13-214) measured in isolation (blue trace). Mixing with equimolar concentration of MtClpP1 (residues 7-200) (green trace) leads to the formation of MtClpP1P2 complexes (purple trace). (C) As in (A) but for MtClpP2 (residues 16-214), with concentrations as indicated. In all panels 0.5 mL of protein at the denoted concentration (monomer) was injected. (D) Peptidase activity assays on a pair of mixtures containing MtClpP2 (residues 13-214) (black bar) or MtClpP2 (residues 16-214) (grey bar) performed in the presence of MtClpP1 (residues 7-200), 4 mM Bz-LL, with 250 µM Suc-LY-AMC used as substrate. In both cases the (monomer) concentration of each of MtClpP1 and MtClpP2 is 1 µM. Error bars correspond to one standard deviation based on three measurements.

**Figure S2.**
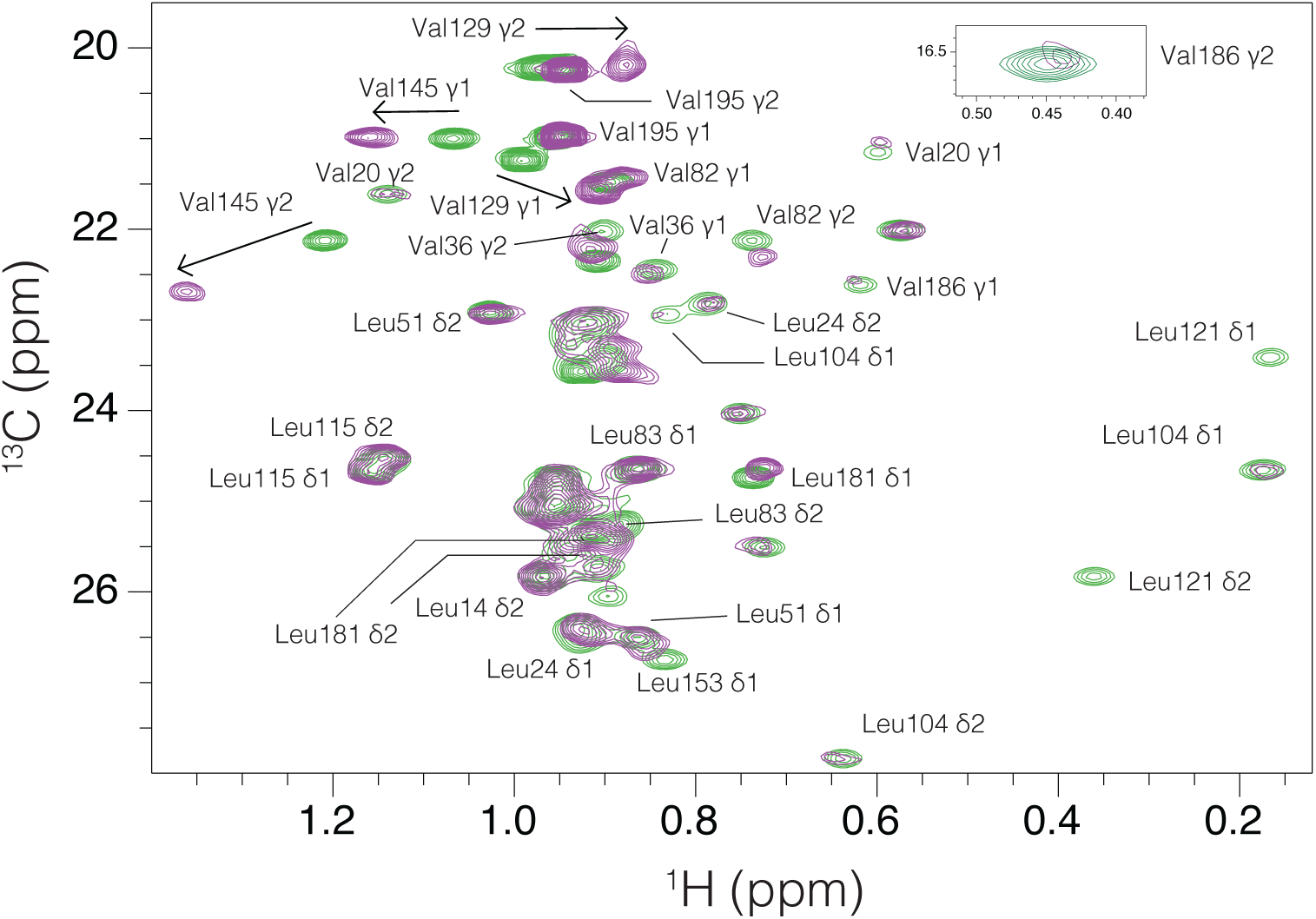
Overlay of the Leu and Val regions of ^1^H-^13^C HMQC correlation maps of ILVM-labeled MtClpP1 (green contour) and a mixture containing 50 µM ILVM-labeled MtClpP1 and 100 µM [U-^2^H] MtClpP2 (purple contours), recorded at 40 °C, 18.8 T, with stereospecific assignments as indicated.

**Figure S3.**
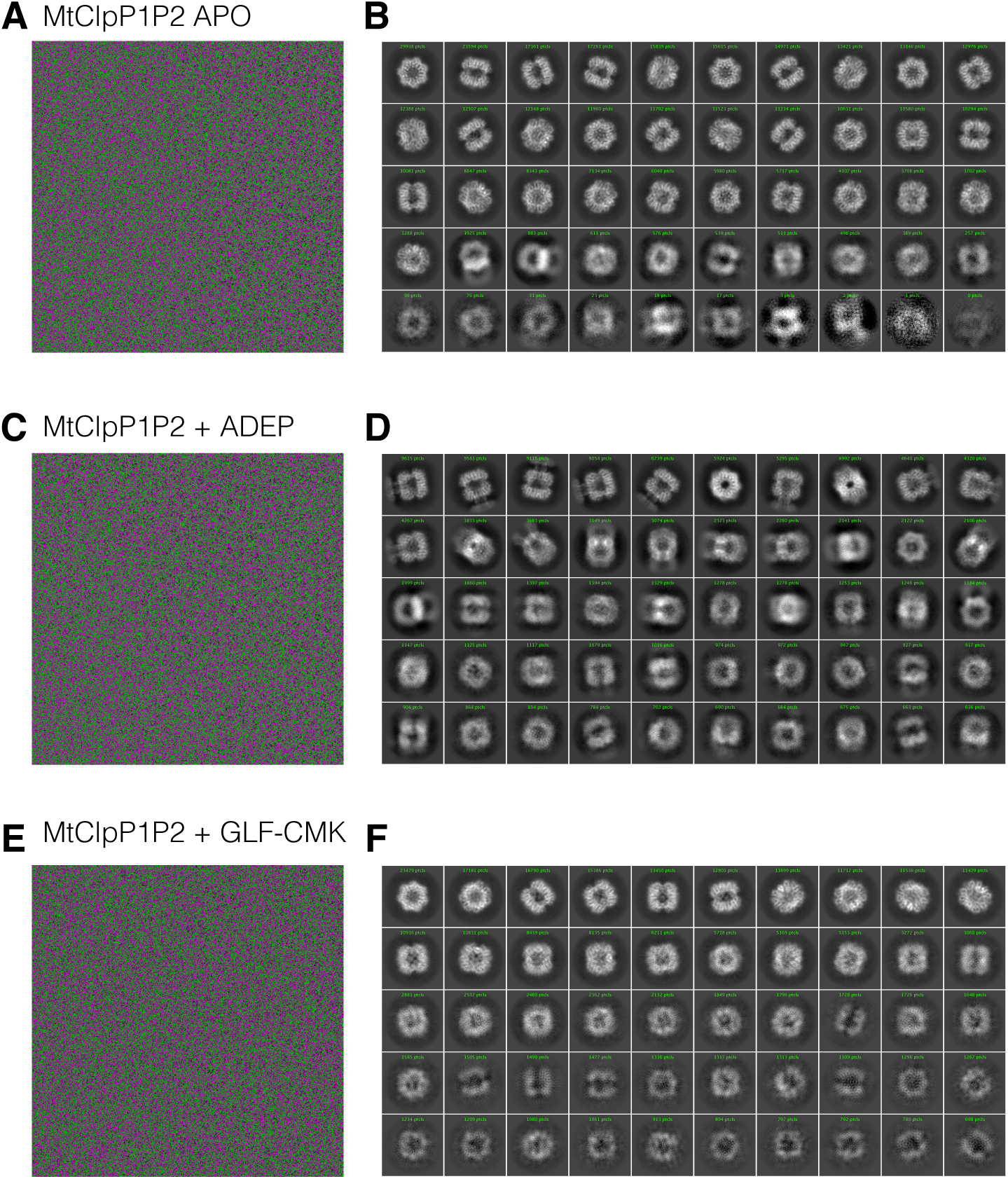
Representative electron micrographs and 2D classes of the MtClpP1P2 complexes in the (A&B) apo, (C&D) ADEP-bound, and (E&F) GLF-CMK bound forms of the enzyme. The number of particles used in each class average is indicted.

**Figure S4.**
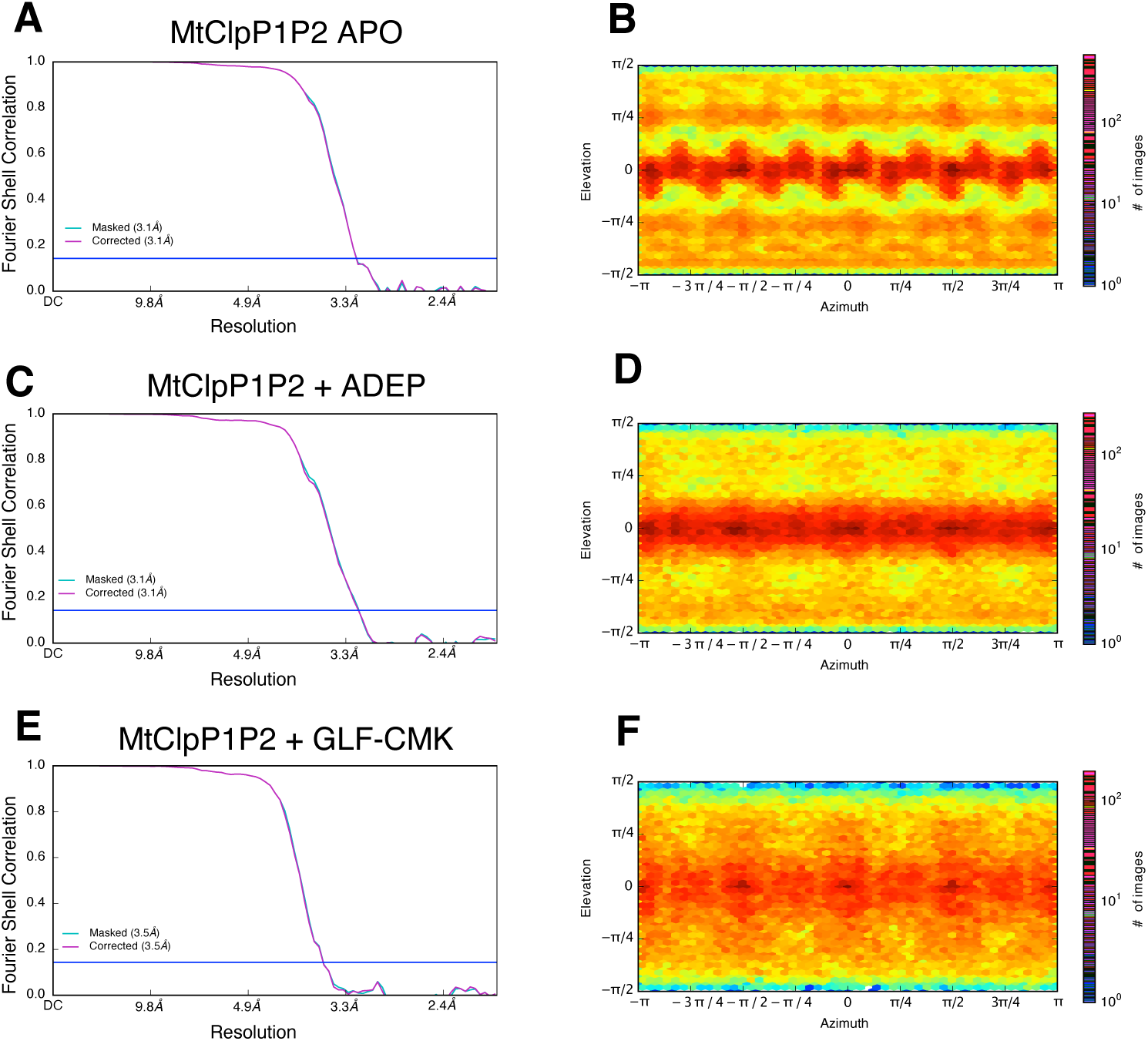
Fourier Shell Correlation (FSC) as a function of spatial resolution and orientation plots for MtClpP1P2 complexes in the (A&B) Apo, (C&D) ADEP-bound, and (E&F) GLF-CMK bound forms. Resolution values reported are for FSC=0.143.

**Figure S5.**
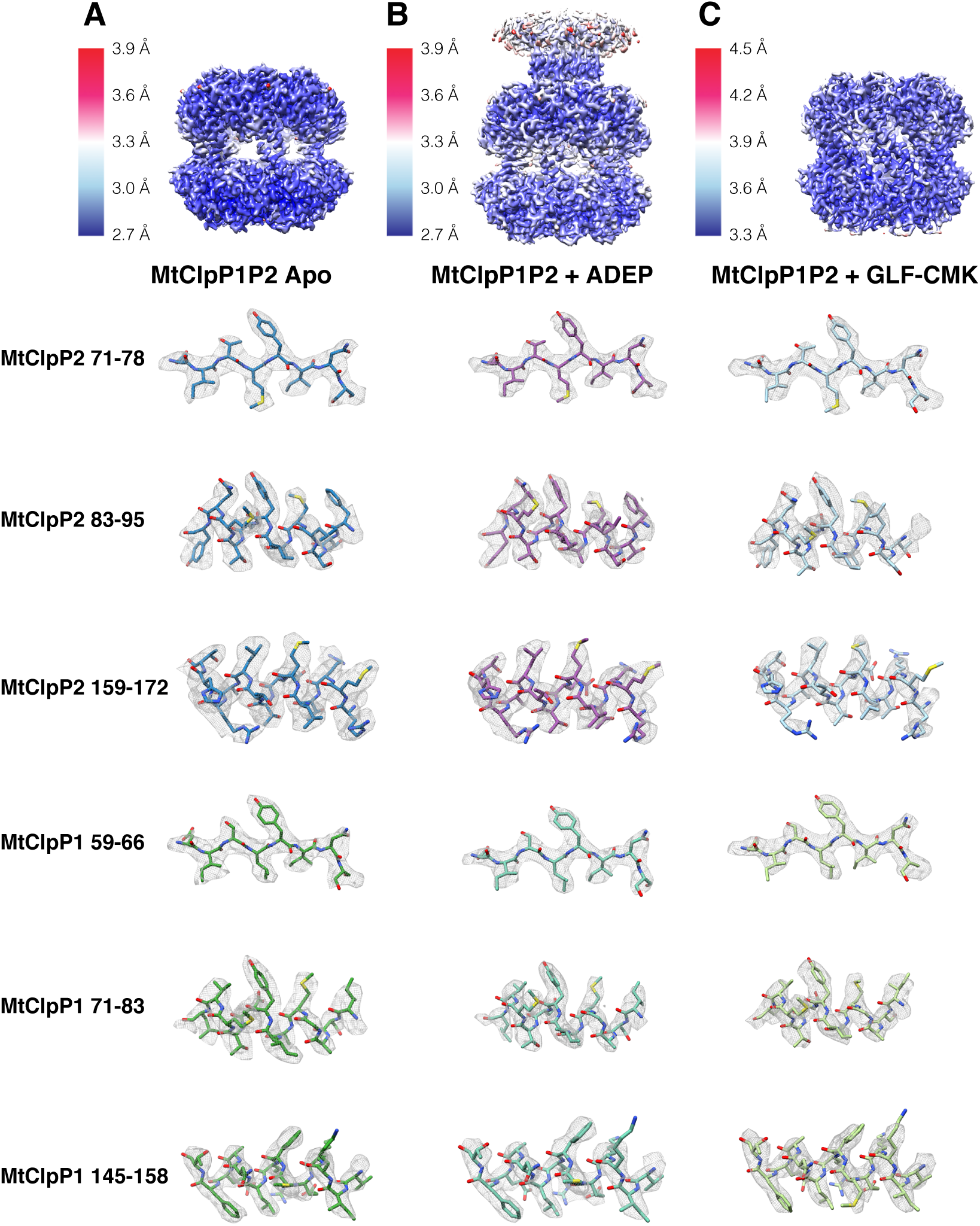
Local resolution maps of MtClpP1P2 complexes in the (A) Apo, (B) ADEP-bound, and (C) GLF-CMK bound forms. Note that in the ADEP-bound state particles are ‘dimers’ of tetradecamers linked via the gates of the P2 ring. Example regions of models built into the experimental cryo-EM maps are also shown (D).

**Figure S6.**
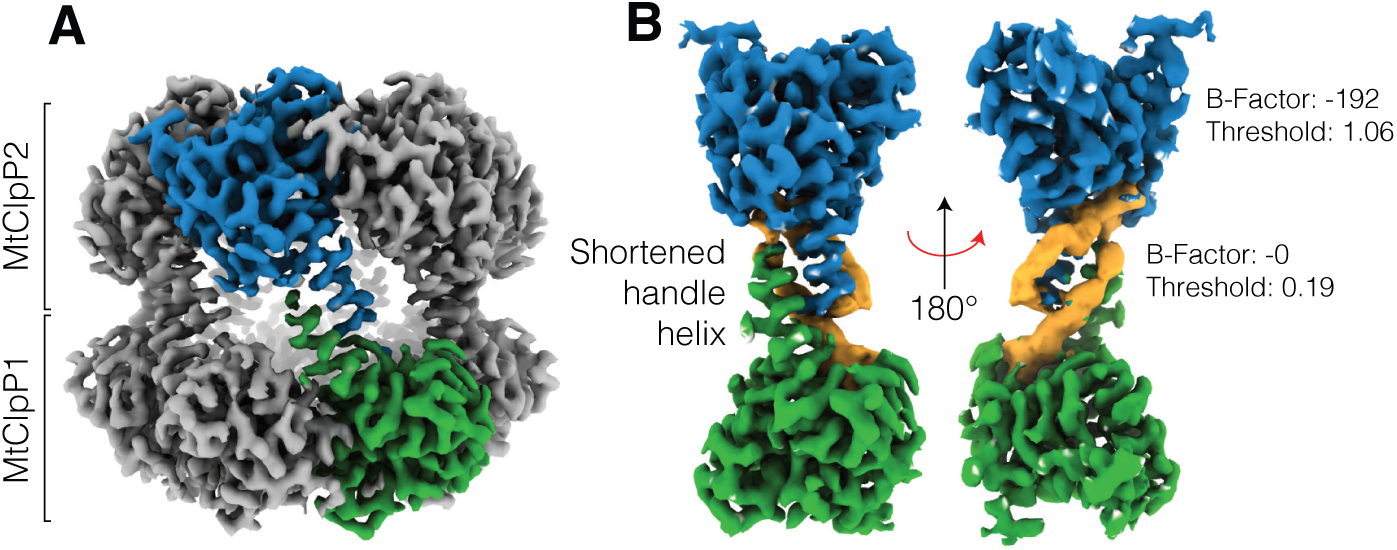
(A) Density map for the apo MtClpP1P2 complex. Single MtClpP1 and MtClpP2 protomers in the complex are coloured green and blue, respectively. (B) Density for a single protomer pair is shown along with an unsharpened map for the flexible handle region.

**Figure S7.**
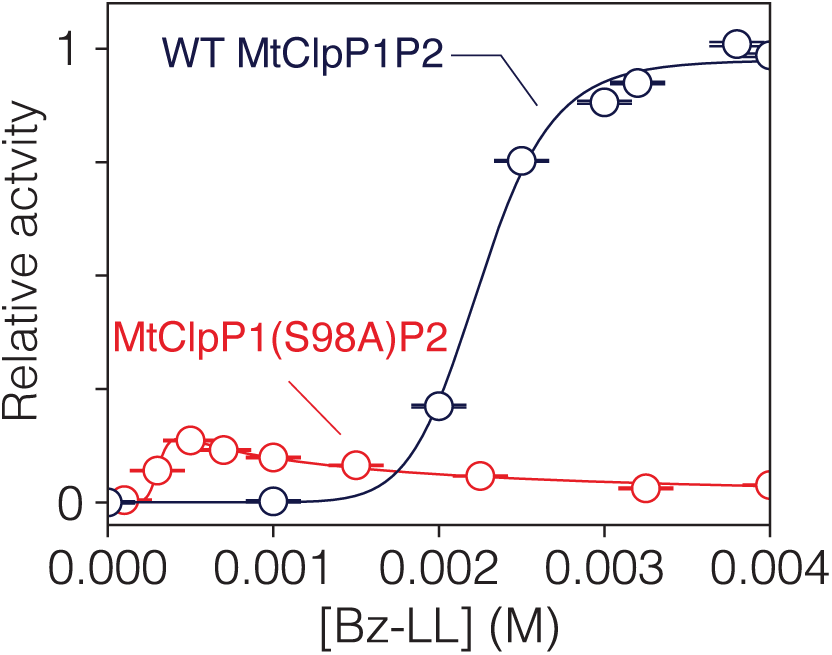
Activity response curves measured as a function of Bz-LL concentration using 250 µM PKM-AMC as substrate for WT MtClpP1P2 (black curve) and MtClpP1(S98A)P2 (red curve). The shift in the shape of the activity curve to lower Bz-LL concentrations for MtClpP1(S98A)P2 is due to the S98A mutation in MtClpP1 (see Discussion).

**Figure S8.**
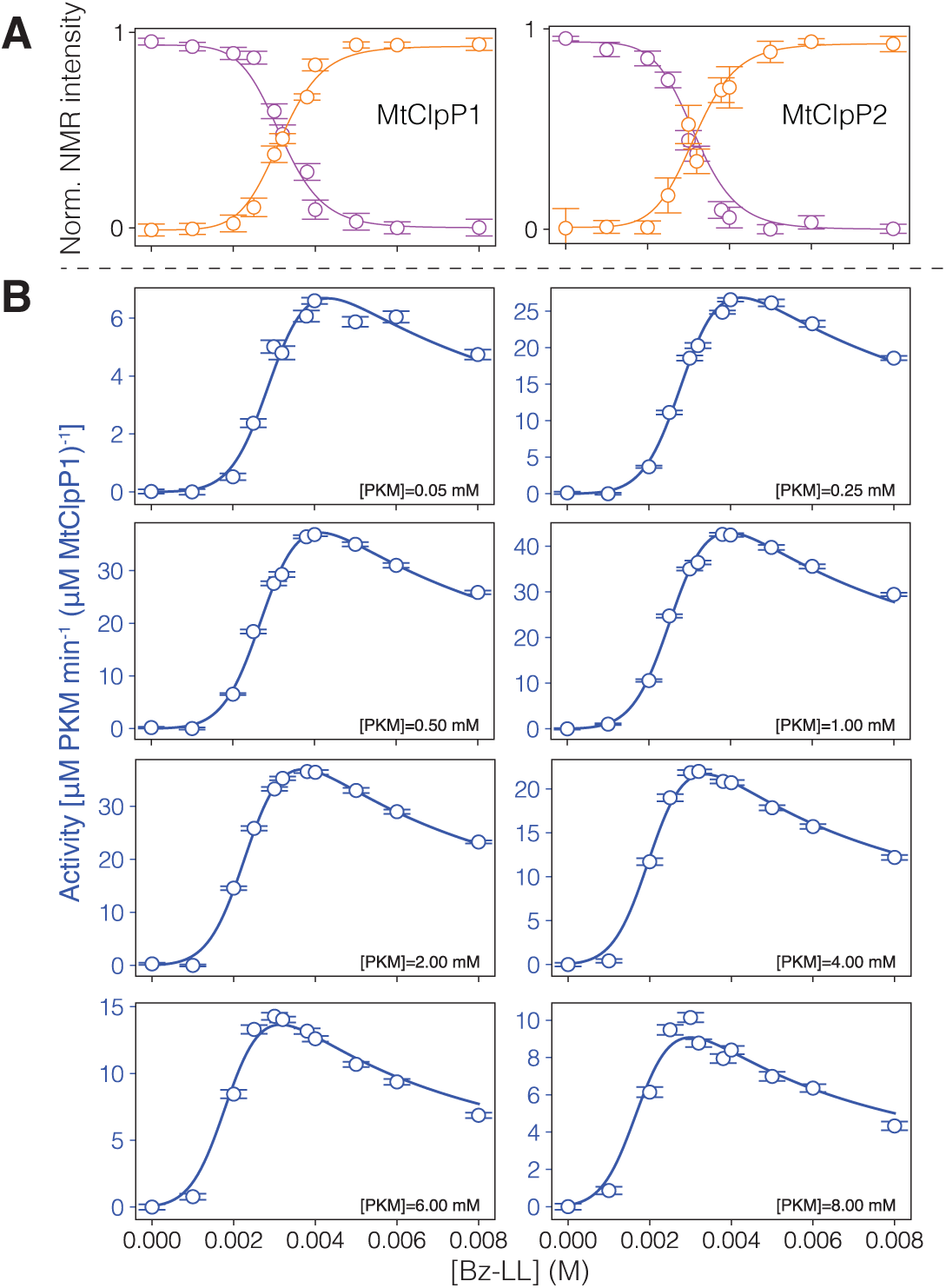
Combined fits of (A) NMR peak intensities and (B) MtClpP1P2 activity as a function of the concentration of Bz-LL. The modified MWC model illustrated in Figure 4D has been used which includes binding of Bz-LL and substrate to the *T* and *R* states of MtClpP1 and MtClpP2 of the complex.

**Figure S9.**
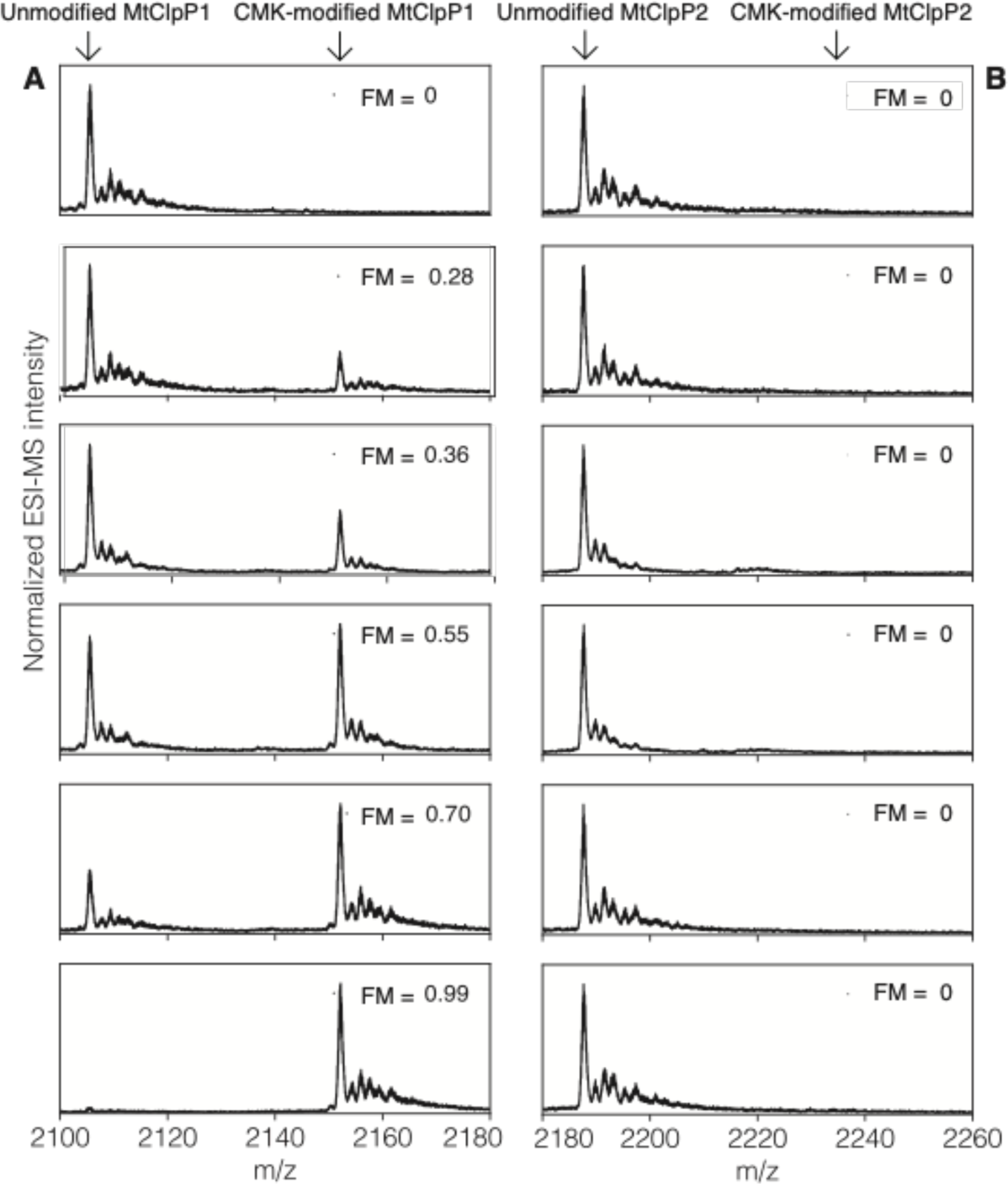
Intact protein mass spectra of each ring of MtClpP1P2 partially modified by GLF-CMK. The 10+ charge state for each of (A) MtClpP1 and (B) MtClpP2 along with expected positions of the unmodified and GLF-CMK modified protein peaks are indicated with arrows. The fraction of active sites modified (FM) is indicated for each panel.

**Figure S10.**
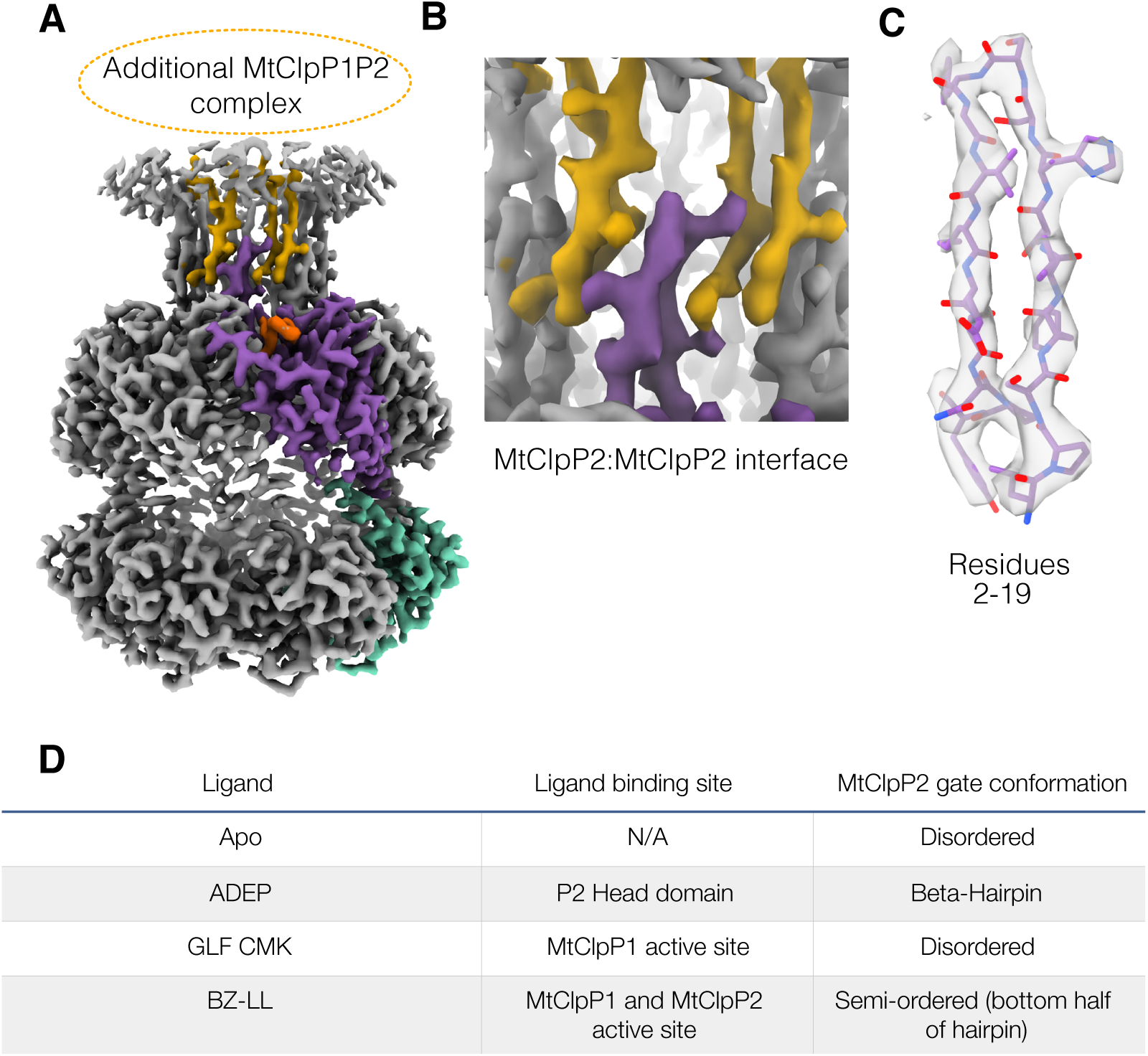
ADEP bound MtClpP2 gates and density. (A) Density map for the MtClpP1P2 complex bound to ADEP. Additional density was present above the MtClpP2 N-terminal gates corresponding to a second MtClpP2 ring that results from dimerization of the complex upon addition of ADEP. Single MtClpP1 and MtClpP2 protomers are coloured teal and purple, respectively. Gate density for two protomers of the second P2 ring are coloured yellow. (B) Magnified view of the MtClpP2:MtClpP2 interface mediated by the N-terminal gates. (C) Model in map fit for a single N-terminal gate. (D) Table summarizing gate conformations in all the states observed for MtClpP1P2.

**Table 1.** Cryo-EM data acquisition, processing, atomic model statistics, and map/model depositions.

A. Cryo-EM data acquisition and image processing.
| Data Collection |  |  |  |
| --- | --- | --- | --- |
| Electron Microscope | Titan Krios |  |  |
| Camera | Falcon 3EC |  |  |
| Voltage (kV) | 300 |  |  |
| Nominal Magnification | 75,000 |  |  |
| Calibrated physical pixel size (Å) | 1.06 |  |  |
| Total exposure (e/Å<sup>2</sup>) | 42.7 |  |  |
| Exposure rate (e/pixel/s) | 0.8 |  |  |
| Number of frames | 30 |  |  |
| Defocus range (µm) | 0.7 to 2.0 |  |  |
| Image Processing |  |  |  |
|  | MtClpP1P2 APO | MtClpP1P2 + ADEP | MtClpP1P2 + GLF-CMK |
| Motion correction software | *cryoSPARC v2* | *cryoSPARC v2* | *cryoSPARC v2* |
| CTF estimation software | *cryoSPARC v2* | *cryoSPARC v2* | *cryoSPARC v2* |
| Particle selection software | *cryoSPARC v2* | *cryoSPARC v2* | *cryoSPARC v2* |
| Micrographs used | 2,092 | 725 | 1645 |
| Particle images selected | 612,408 | 366,129 | 257,060 |
| 3D map classification and refinement software | *cryoSPARC v2* | *cryoSPARC v2* | *cryoSPARC v2* |

B. Map and model statistics.
| EM maps | MtClpP1P2 APO | MtClpP1P2 + ADEP | MtClpP1P2 + GLF-CMK |
| --- | --- | --- | --- |
| Particle images contributing to maps | 373,064 | 192,430 | 143,748 |
| Applied symmetry | C7 | C7 | C7 |
| Applied B-factor (Å<sup>2</sup>) | -191.8 | -168.6 | -217.4 |
| Global resolution (FSC = 0.143, Å) | 3.1 | 3.1 | 3.5 |
| Model Building |  |  |  |
| Modeling software | Coot, Phenix, Rosetta |  |  |
| Number of residues | 2,513 | 2,618 | 2,506 |
| RMS bond length (Å) | 0.0196 | 0.0202 | 0.0049 |
| RMS bond angle (°) | 1.84 | 1.70 | 1.08 |
| Ramachandran outliers (%) | 0.85 | 0.54 | 0 |
| Ramachandran favoured (%) | 95.77 | 96.22 | 98.31 |
| Rotamer outliers | 0 | 0 | 0 |
| C-beta deviations | 0 | 0 | 0 |
| Clashscore | 0.26 | 2.66 | 1.16 |
| MolProbity score | 0.89 | 1.31 | 0.83 |
| EMRinger score | 4.3 | 3.7 | 2.41 |
| Ligand | N/A | ADEP-7 | GLF-CMK |

C. Residues excluded in atomic models.
| ClpX Protomer | MtClpP1P2 APO | MtClpP1P2 + ADEP | MtClpP1P2 + GLF-CMK |
| --- | --- | --- | --- |
| ClpP1 | 1-15,193-200 | 1-15,193-200 | 1-15,193-200 |
| ClpP2 | 1-30, 211-214 | 1-15, 211-214 | 1-31, 211-214 |

D. Deposited maps and associated coordinate files.
| Maps | EMDB code | Associated PDB ID |
| --- | --- | --- |
| MtClpP1P2 APO | EMD-XXXX | XXXX |
| MtClpP1P2 + ADEP | EMD-XXXX | XXXX |
| MtClpP1P2 + GLF-CMK | EMD-XXXX | XXXX |

